# Threats to Nature’s contributions to people provided by terrestrial vertebrates across Europe

**DOI:** 10.1101/2025.10.25.684519

**Authors:** Louise O’Connor, Francesca Cosentino, Servane Demarquet, Pierre Gaüzère, Tom Hackbarth, Luigi Maiorano, Chiara Mancino, Julien Renaud, Sara Si-Moussi, Peter H. Verburg, Wilfried Thuiller

## Abstract

**Aim.:** Species and ecosystem processes offer essential benefits to people, known as Nature’s Contributions to People (NCP). However, we still lack a comprehensive understanding of NCP provided by terrestrial vertebrates on a large scale, and of the threats they face. To bridge this gap, we built a comprehensive dataset that documents the NCP provided by terrestrial vertebrate species in Europe, and analysed the conservation status and threats to NCP provider species.

**Location.:** Europe

**Methods.:** We synthesised existing literature on NCP associated with European terrestrial vertebrates, and leveraged ecological traits and trophic interactions from previously established datasets. We identified 15 NCP (10 regulating NCP and 5 non-material NCP), with 860 species providing at least one NCP (out of 1,168 vertebrate species considered in total). Then, we harnessed species distribution data and a novel European land system map to create species-mediated NCP maps across Europe at a 1km² resolution, including societal demand for each NCP.

**Results.:** We found that i) for each NCP, at least 25% of NCP provider species are assessed as threatened with extinction; ii) NCP multifunctionality is lowest in high-intensity land systems; and iii) direct exploitation and agricultural intensification are major threats to species-mediated NCP, impacting both non-material and regulating NCP provider species.

**Main conclusions.:** Protecting threatened NCP provider species, and reducing direct exploitation are key to maintain regulating and non-material NCP. Our results suggest that de-intensifying agricultural practices, through maintaining heterogeneous mosaic landscapes and promoting diversified practices, could increase NCP multifunctionality. Our work enables a comprehensive understanding of NCP provided by terrestrial vertebrates in Europe, their biogeography, and the threats they face, which can in turn inform spatial conservation planning to improve the conservation of both biodiversity and NCP.

## 1. Introduction

Knowledge of the functional links between species and nature’s contributions to people (NCP) is crucial for effective conservation policy. The concept of Nature’s Contributions to People (NCP) extends and refines the earlier framework of ecosystem services, which was criticised for commodifying nature and overlooking non-material values. The NCP concept offers a more inclusive and broader approach to understanding the relationship between biodiversity and human well-being (Díaz et al. 2018). NCP describe the various ways in which ecosystems sustain human life, including benefits that are material (food, energy, timber), non-material (recreation, inspiration) and regulating (climate regulation, pollination, erosion control) (Díaz et al. 2018). The Intergovernmental Science-Policy Platform on Biodiversity and Ecosystem Services (IPBES) recognizes 18 distinct categories of NCP, encompassing a wide range of material, regulating, and non-material contributions (Table 1). The ecosystem services framework originated in plant functional ecology (Diaz et al., 2007), and many studies seeking to describe and understand species contributions to people have focused on the contributions by plants and the above-and below-ground communities they interact with, such as pollinators (Lavorel 2013; Harrison et al. 2014; Pironon et al. 2024). By contrast, the role of vertebrates in providing NCP remains comparatively understudied and has so far focused either on specific taxonomic groups (e.g. amphibians: Hocking & Babbitt 2014; birds: Smith et al. 2022; carnivores: Frank 2024), particular ecosystems (e.g. alpine areas: Lavorel et al. 2020; agroecosystems: Civantos et al. 2012), or individual countries (e.g. Switzerland: Rey et al. 2023; Brazil: Vale et al. 2023).

**Table 1.** Overview of NCP considered in the study, data sources, and link to broader NCP categories.

| Broad NCP categories from (Diaz et al., 2021) | Vertebrates' contributions to people | Sources |
| --- | --- | --- |
| Habitat creation and maintenance (regulating) | Keystone species and ecosystem engineers | Apex predators were identified from the metaweb, and ecosystem engineers from the literature |
| Regulation of detrimental organisms and biological processes (regulating) | Agricultural pest control | Predators of invertebrate and rodent pests were extracted from Civantos et al. (2012), and complemented with the metaweb. |
| Regulation of detrimental organisms and biological processes (regulating) | Mosquito control (potential disease vectors) | Mosquito predators were identified from the metaweb. |
| Regulation of detrimental organisms and biological processes (regulating) | Tick host biocontrol (disease control) | Predators of ungulates and rodents, both important reservoirs of tick-borne diseases, were identified from the metaweb. |
| Regulation of detrimental organisms and biological processes (regulating) | Pine processionary moth control | Bird predators were extracted from Barbaro and Battisti (2011); bat predators were identified from the metaweb. |
| Regulation of detrimental organisms and biological processes (regulating) | Carrion elimination (related to disease control and nutrient cycling) | Frequent or obligatory scavengers were identified from the metaweb, and complemented with the global database of scavengers (Sebastián-Gonzalez et al., 2021). |
| Regulation of detrimental organisms and biological processes (regulating) | Regulation of an invasive alien species: the Asian hornet | Its predators are: *Pernis apivorus* (Macia et al., 2019) and *Merops apiaster* (Onofre et al., 2023). |
| Regulation of detrimental organisms and biological processes (regulating) | Bark beetle control | Bark beetle predators were extracted from Wegensteiner et al. (2015). |
| Pollination and dispersal of seeds and other propagules; Habitat creation and maintenance (regulating) | Seed dispersal (related to vegetation regeneration and carbon storage) | Frugivorous species (endozoochory) were identified from the metaweb; other seed dispersers (ecto- and syn-zoochory) were extracted from Gomez et al. (2019) and Mendes et al. (2024). |
| Pollination and dispersal of seeds and other propagules (regulating) | Pollination (agriculture) | Bird pollinators were extracted from (da Silva et al., 2014). |
| Supporting identities (non-material NCP) | Species of cultural importance (SCI) | SCI were identified from the Birds and Habitats directive; and Wikipedia page visitation rates. |
| Supporting identities; Physical and psychological experiences | Game | Game species were extracted from Schulp et al. (2014). |
| (non-material NCP) |  |  |
| Maintenance of options;<br>supporting identities (non-material NCP) | Evolutionary heritage | EDGE species were extracted from EDGE Lists (2018). |
| Learning and inspiration;<br>Physical and<br>psychological experiences<br>(non-material NCP) | Nature tourism and<br>wildlife watching | Popular wildlife watching species were extracted from touristic webpages (Table S1) and density of observations (GBIF) across species' European range. |
| Learning and inspiration;<br>Physical and<br>psychological experiences<br>(non-material NCP) | Birdsong | Songbirds were identified based on taxonomy. |

There is growing evidence that terrestrial vertebrates deliver a wide array of NCP at continental and global scales, including many regulating and non-material NCP (Ceau_u et al. 2021; Chaplin-Kremer et al., 2025). Vertebrates provide many regulating contributions to people via their trophic role and biotic interactions (Antunes et al. 2024). Pollination by birds and bats supports reproductive success in flowering plant species (da Silva et al., 2014; Ramirez-Francel et al., 2021); seed dispersal by frugivorous vertebrates, large mammals transporting seeds on their hooves or fur, or by scatter-hoarders, underpins plants regeneration and indirectly contributes to carbon sequestration (Mendes et al., 2024). In agricultural landscapes, predator species regulate populations of insect and rodent pests (Civantos et al., 2012); and in forests, insectivorous predators regulate pests such as pine processionary moths and bark beetles. Emerging evidence highlights the substantial economic and health benefits brought by the natural control of pests by their predators. For example, the collapse of insectivorous bat populations in the United States (as a consequence of an invasive fungus disease) resulted in a 31% increase in pesticide use, leading to negative impacts on human health and a rise in infant mortality (Frank, 2024). Terrestrial vertebrates may also contribute to the regulation of invasive alien species through predation, thereby mitigating the substantial ecological and economic impacts associated with biological invasions (Diagne et al., 2021). In addition to regulating services, terrestrial vertebrates also provide non-material contributions to people. Birdsong has been associated with psychological well-being and mental health benefits that appear to be distinct from the benefits of simply being in nature (Hammoud et al., 2022; Stobbe et al., 2022). Numerous charismatic bird and mammal species underpin nature-based tourism and wildlife watching, and many vertebrate species are of high cultural importance to a range of European people and cultures (Reyes-García et al. 2023).

NCP value is higher, and more stable, when there is a higher diversity of species that provide it (Harrison et al., 2014; Gutierrez-Arellano and Mulligan, 2018; Ceausu et al., 2021). For example, a high richness of predators can stabilise local NCP capacity by controlling pest populations more effectively and consistently through time, compared with a single predator species (Ostfeld & Holt, 2004). Further, different species often occupy different functional and trophic niches and complement each other’s roles in providing NCP. For example, a community comprising both diurnal insectivorous birds and nocturnal insectivorous bats will ensure a more effective regulation of insect pests, as these predators will feed at different times on pest species. Similarly, different species may regulate different life stages of a pest, with amphibians predating on the larval life stage of the mosquito, while bats predate on the adult mosquito; and the caterpillar of the pine processionary moth is eaten by insectivorous birds while the moth is eaten by insectivorous bats. Species richness and functional diversity thus increase the local NCP value and stability (Harrison et al., 2014). Species richness and functional diversity act as insurance for ecosystem functioning, making NCP provided by vertebrates more resilient to disturbances (Yachi and Loreau 1999). Furthermore, diverse vertebrate communities will likely increase multifunctionality across NCP: e.g. a community composed of pollinators, seed-dispersing frugivores, insectivores, and scavengers will support plant regeneration, crop productivity, and carrion removal. Yet, global defaunation is now recognized as a major threat to the continued delivery of vertebrate-mediated NCP, because of the loss or decline of functionally unique species (Chaplin-Kramer et al. 2025; Fricke et al. 2022). For example, seed dispersal is one of the most widespread ecosystem functions provided by vertebrates, yet this function is increasingly threatened by habitat loss, fragmentation, invasive species, and direct exploitation (Mendes et al., 2024; Fricke et al. 2022).

Despite advances in exploring vertebrates’ contributions to people, a comprehensive and spatially explicit synthesis at a continental scale is thus far still lacking. As conservation policies increasingly aim to safeguard both biodiversity and nature’s contributions to people, a broader understanding on the biogeography and vulnerability of vertebrates’ contributions to people is critical, for improving our understanding of the potential synergies and trade-offs between safeguarding biodiversity and sustaining NCP, and for guiding the prioritisation of species and areas in need of conservation or restoration (Ceau_u et al. 2021; O’Connor et al., 2021; Chaplin-Kramer et al. 2025). Here, we provide a continental-wide assessment of the linkages between all terrestrial vertebrate species and fifteen different NCP in Europe. Our objectives were to 1) compile a comprehensive database linking European terrestrial vertebrate species to the regulating and non-material NCP they provide; 2) quantify the conservation status and major threats to provider species ; 3) examine how land use intensity and land cover interact to shape local NCP multifunctionality across Europe; 4) explore spatial patterns of NCP supply. To address these goals, we created a dataset summarizing the NCP provided by 1,168 European vertebrate species, by integrating published literature and datasets on NCP provided by vertebrates, their functional traits, trophic interactions. Combining this knowledge with species distributions, we quantified and mapped each NCP and further examined spatial patterns of NCP capacity. We also mapped the societal demand for each NCP across European land systems (Sandström et al., 2023) and combined it with NCP capacity in order to create NCP supply maps and highlight areas with both high capacity and high demand for the NCP (Verhagen et al., 2017). This is a crucial step if we are to identify conservation priorities that provide benefits where they are most needed. We hypothesized that most NCP providers are not threatened with extinction (Rey et al., 2024); that high land use intensity decreases NCP supply; and that spatial patterns vary between NCP and European regions.

## 2. Methods

### Overview

We compiled a spatial database on the NCP provided by European vertebrate species, and described the biogeography and vulnerability of vertebrates’ contributions to people in Europe. We first identified ten regulating NCP and five non-material NCP linked to European terrestrial vertebrates and created a binary matrix summarising the links between each species and NCP. We used published datasets on NCP provided by vertebrate species (Table 1), as well as data on species’ functional traits, diets, and trophic interactions (Thuiller et al. 2015; Maiorano et al. 2020). The species pool we considered covered all (native and breeding) terrestrial vertebrate species known to occur in Europe: this includes 1,168 terrestrial vertebrate species in total: 292 mammals, 528 birds, 247 reptiles and 101 amphibians (Fig. 1). Second, we mapped species-mediated NCP across Europe at a resolution of 1 km, using species distributions data (Si-Moussi and Thuiller, 2024) and a novel European land system map (Dou et al. 2021). We further integrated capacity and demand to map NCP supply (sensu Verhagen et al. 2017; O’Connor et al. 2021) by combining provider richness and by accounting for the extent to which there is societal demand for each NCP in different land systems (defined in Table S2). Third, we analysed current threats to species-mediated NCP in Europe by describing: i) how land use intensity is related to NCP provider richness ; ii) the conservation status of species providing NCP; and iii) which anthropogenic threats are most impactful to each of the species-mediated NCP.

**Figure 1:**
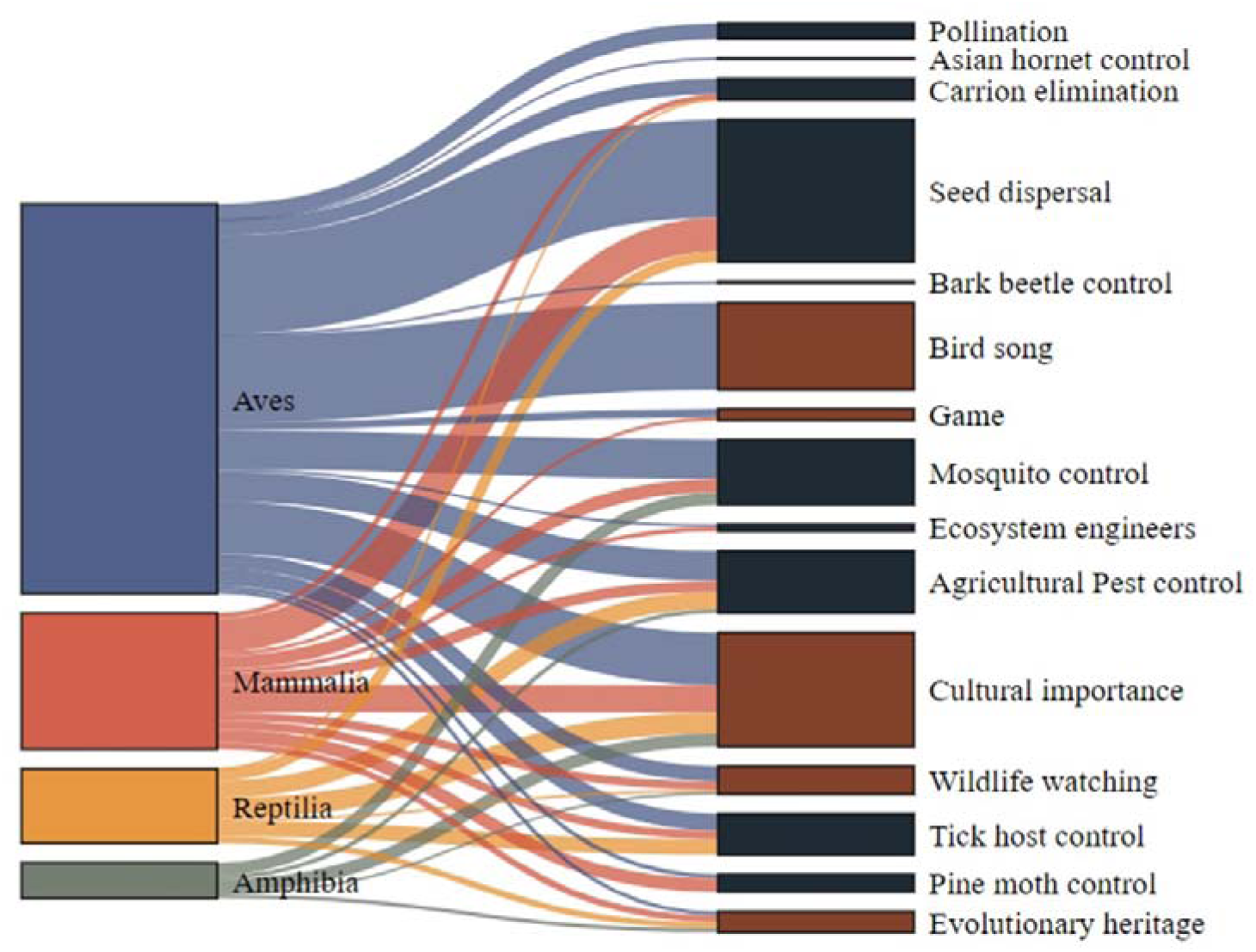
Relationships between vertebrate provider species, grouped by taxonomic class, and each of the 15 NCP. Links in the Sankey diagram represent the connections between species and NCP provision and they are coloured by the taxonomy of the provider species. Link width is proportional to the number of species providing the NCP in each taxonomic class. Non-material NCP are shown in brown, and regulating NCP in dark blue.

### 2.1. Building the database of species-mediated NCP

#### 2.1.1. Selecting relevant NCP

We initially searched the literature on NCP provided by terrestrial vertebrates, specifically in the context of Europe, using keywords “services”, “NCP” together with “vertebrates”, “birds”, “mammals”, “reptiles”, “amphibians”. This led to an initial list of 22 different NCP. We then Then, we refined our initial list and excluded 1) NCP for which we lacked sufficient evidence or data; 2) those that were highly context specific and unlikely to be relevant across all of Europe (e.g. frog consumption); and 3) those that are associated with negative trade-offs (e.g., large wild animals may play a role in mitigating climate change but whose functional role may be associated with trade-offs or conflicts (Malhi et al., 2022)). Thus, for the analyses, we retained a set of fifteen NCP that are positive, broadly relevant across Europe, and with sufficient supporting evidence. The final selection covered ten regulating NCP (keystones and ecosystem engineers, agricultural pest control, mosquito control, tick host control, pine moth control, bark beetle control, invasive Asian hornet control, carrion elimination, seed dispersal, pollination) (Fig. S1) and five non-material NCP (cultural importance, evolutionary heritage, nature tourism and wildlife watching, birdsong, and game) (Fig. S2).

#### 2.1.2. Identifying provider species for regulating NCP

##### Agricultural pest biocontrol

Control of agricultural pests by natural predators is an essential NCP provided by vertebrate species which cannot be effectively replaced with pesticide use (Frank 2024). We extracted the species in the dataset built by (Civantos et al. 2012), which provides the list of species that feed on rodent and/or invertebrate species that are considered pests for agriculture in Europe. For rodent pest control, we additionally extracted natural predators of two major rodent pest species in European agricultural landscapes: *Microtus arvalis* and *Apodemus sylvaticus*. This led to an additional 89 predators of rodent pests compared to the list of rodent pest control providers identified in Civantos et al. (2012). In total, there were 195 species that provide agricultural pest control.

##### Mosquito biocontrol

Mosquitoes are vectors for diseases which may become widespread in Europe due to climate change (Semenza and Paz, 2021). Insectivorous bats and birds with aerial foraging space act as predators of the adult life stage of mosquitoes (Puig-Montserrat et al. 2020), while amphibians that feed primarily on aquatic invertebrates regulate its larval life stage (Hocking and Babbitt 2014). To identify species that predate on mosquitoes, we used the Tetra-EU metaweb of trophic interactions between all vertebrate species in Europe (Maiorano et al. 2020) and other diet categories. This metaweb distinguishes obligate or frequent interactions from occasional feeding interactions. Here, we only considered the obligate feeding interactions, because these represent significant and frequent energy flow, and are more likely to underpin mosquito control. We thus assumed that occasional interactions are not representative of a species’ feeding ecology, and therefore only make marginal contributions to NCP supply. Combining the diet and the foraging space of species, we identified a total of 202 species (consisting of aerial feeding bats and birds, and amphibians) that can regulate populations of mosquitoes in Europe.

##### Biocontrol of the invasive Asian hornet

The honey buzzard (*Pernis apivorus)* is a proven predator of the invasive Asian hornet nests (Macià et al. 2019) and is likely an effective agent for Asian hornet biocontrol because of its behaviour of destroying the nests. There is also evidence for the European bee-eater (*Merops apiaster*) to act as a biocontrol agent for the Asian hornet (Onofre et al., 2023). These are the only two known predators of the invasive Asian hornet in Europe.

##### Pine processionary moth control

The pine processionary moth has been listed as a species of concern to human health (de Boer and Harvey 2020). One of the most effective means to control their spread is biocontrol by predators. Cuckoos (*Clamator glandarius* and *Cuculus canorus*), Eurasian hoopoe (*Upupa epops*), and birds of the tit family (*Paridae*), are known predators of the caterpillar life stage (Barbaro and Battisti 2011). In addition, insectivorous bats are predators of the adult lifestage: we identified insectivorous bats that can regulate pine moths from the metaweb (Ramirez-Francel et al., 2021). In total, we identified 55 species that can provide biocontrol of the pine processionary moth.

##### Regulation of tick-borne diseases

Ticks are widespread across Europe, and large, unchecked populations of tick hosts, especially ungulates and rodents, are important reservoirs for tick borne pathogens (e.g. Borrelia). Predators of rodents and of ungulates can help control the spread of tick-borne disease and protect human health (Černý et al. 2020; Hofmeester et al. 2017). We extracted species that are obligate predators of rodents and of ungulates from the Tetra-EU metaweb (Maiorano et al., 2020), described above. Using the metaweb data, we identified 128 species that are obligate predators of rodents and/or ungulates, and therefore that can provide tick host biocontrol.

##### Carrion elimination

The removal of carcasses by scavengers reduces the risk of disease transmission and contributes to nutrient cycling in the ecosystem (Mateo-Tomás et al. 2017; Vicente and VerCauteren 2019; Probst et al. 2019). Using the diet categories in the metaweb database (Maiorano et al., 2020), we identified 52 species of scavengers that provide carrion elimination. We further included terrestrial vertebrate scavengers documented in the global scavenger database (Sebastián-Gonzalez et al., 2021). Among the 177 listed in the database, we extracted 55 that occur in Europe. However, this database included species that would normally not feed on carrion as part of their diet: *Dryocopus martius, Cyanistes caeruleus, Genetta genetta, Herpestes ichneumon, Lynx lynx, Bison bonasus, Martes martes, Parus major, Dendrocopos major, Lynx pardinus, Hystrix indica, Sciurus vulgaris, Lanius collurio, Mustela erminea,* and *Felis silvestris*. We excluded these species and retained the remaining 40 species documented in the database.

##### Keystone species and ecosystem engineers

Some large animals play a role in creating and maintaining habitats for other species (Malhi et al., 2022). Top predators often play a keystone role due to their top-down stabilizing effects on communities (Estes et al., 2011; Ripple et al., 2014). We identified top predators from the European metaweb. We defined apex predators as species that occupy the highest trophic levels in the metaweb (trophic level > 2), with no predators in their adult life stage and at least 5 vertebrate prey, and that feed on medium to large-sized prey (Estes et al., 2011). Following these criteria, apex predators in the metaweb were: *Canis lupus, Lynx lynx, Ursus arctos, Gulo gulo, Bubo bubo, Strix nebulosa, Aquila chrysaetos*. A number of species also create and shape habitats for other species and are known as ecosystem engineers. Ecosystem engineers among European vertebrates are: *Castor fiber* (Rosell et al., 2005), *Marmota marmota* (Ballova et al., 2019), *Meles meles* (Kurek et al., 2022), *Sus scrofa* (Barrios-Garcia and Ballari, 2012) and *Cervus elaphus* (Müller et al., 2017).

##### Seed dispersal

Seed dispersal is key to the regeneration of vegetation (especially shrubs and forests) and indirectly supports carbon sequestration. Vertebrates are essential seed dispersers (Sandor, Elphick, and Tingley 2022; Fricke et al. 2022). We considered three different processes of seed dispersal: 1) endo-zoochory by obligate frugivore species, 2) synzoochory by all species with a scatter-hoarding behaviour (e.g. the European jay), known to be highly effective for seed dispersal, and 3) ecto-zoochory. We identified obligate frugivorous species that disperse seeds via endozoochory from the metaweb (Maiorano et al., 2020); scatter-hoarding seed dispersers from Gomez et al., 2019; and all other European vertebrates known to disperse seed via endozoochory, synzoochory or ectozoochory from Mendes et al., 2024. In total, there are 445 species of vertebrate that are known seed dispersers in Europe.

##### Pollination

Pollination is linked to both food supply in pollinator-dependent croplands and to the long term survival of flowering plants through reproduction. Among the terrestrial vertebrates considered here, a few European bird species are occasional pollinators. We extracted 49 species of bird pollinators from the dataset built by (da Silva et al. 2014). Even if occasional, their role as pollinators can be complementary to pollinating insects as they can pollinate over larger distances.

##### Bark beetle control

Bark beetles can cause extensive damage to forests, and their predators, including woodpeckers, can help reduce bark beetle populations. We extracted the list of natural predators of bark beetles from Wegensteiner et al. (2015). In total, four species of woodpeckers occurring in Europe are known to be natural predators of bark beetles.

#### 2.1.3. Identifying provider species for non-material NCP

##### Culturally valued species

In total, we identified 356 terrestrial vertebrate species that are culturally important and valued as part of European natural heritage. We used three different criteria to identify species of cultural importance:

1. Culturally valued species can be identified through their inclusion in the Annexes of the Birds and Habitats Directives, which reflect Europe’s natural heritage and a recognition of species’ ecological, aesthetic or societal importance. We considered terrestrial vertebrate species listed in the Annexes II and IV of the Habitats directive and the Annex I of the Birds directive as culturally significant.
2. We additionally extracted the list of species known to be culturally important in different European socio-cultural groups such as ethnic groups and indigenous people (such as the reindeer in Sami culture), from the dataset by Reyes-Garcia et al., 2023.
3. We additionally identified the top species that are most frequently searched for in Wikipedia, as a proxy for their cultural importance, following previous studies (Roll et al., 2016). We extracted page visitations over a period of 8 years (January 2016 to December 2024) using the pageviews R package. We identified the most visited species based on the inflexion point in the number of page views using the R package inflection which detects the point at which the visitations data changes most sharply. This led to identifying 32 species with visitation rates above the inflection point (Fig. S4).

##### Nature-based tourism and wildlife watching

In total, we identified 55 species of terrestrial vertebrates that are important for nature-based tourism. These consisted of:

1. 44 terrestrial vertebrate species frequently mentioned on touristic websites (Table S1). We used a search engine with the keywords “Wildlife Tourism Europe” and then noted the species mentioned on the websites of parks and wildlife guides.
2. 23 species with a very high number of observations given their small range size. We computed the ratio between the number of observations per species in GBIF and their European range size (km²). We then identified top species with particularly high values of observation density per km² relative to others. To do so, we applied the elbow point detection method using the uik() function from the R package inflection, to detect the point at which the (sorted) observation density data change most sharply. We used the identified inflection point to extract the subset of top-ranking species. This approach provides an objective criterion to define a break in the distribution without requiring arbitrary thresholds. This led to extracting the top 23 species with observation density values above the inflection point (Fig. S5).

##### Wild foods

We extracted 35 terrestrial vertebrate game species that are important for hunting in Europe, from the dataset of wild foods built by (Schulp, Thuiller, and Verburg 2014). Throughout Europe, these species support identities, provide food as well as opportunities for physical experiences in nature.

##### Evolutionary heritage

Evolutionary heritage reflects the unique evolutionary history of a species. Species with a high EDGE index are both globally endangered and evolutionarily distinct, and possess unique traits shared by no other species. Preserving EDGE species therefore safeguards not only species richness but also the breadth and depth of the evolutionary tree of life, and is especially important for the “maintenance of options” dimension of NCP (Diaz et al., 2018).

We extracted the 65 European terrestrial vertebrate species listed as Evolutionarily Distinct and Globally Endangered in the EDGE database (“EDGE Lists” 2018; Isaac et al. 2007).

##### Birdsong

Exposure to birdsong has been shown to benefit mental and psychological well-being and has been shown to alleviate anxiety, stress, and paranoia (Hammoud et al., 2022; Stobbe et al., 2022). These benefits appear to be distinct from the general positive effects of being in nature, suggesting that birdsong plays a unique role in supporting mental health. In Europe, most birds from the *Passeriformes* order (such as thrushes, nightingales, robins, finches, warblers) are known for their melodious singing and were included in our selection, with the exceptions of corvids and sparrows, which we excluded due to their harsh vocalisations. In addition to passerines, we also included a set of bird species with melodic and aesthetic vocalisations. These included certain species of loons, sandpipers, doves, owls, woodpeckers, swifts and swallows, as well as a number of other species with pleasant or culturally valued vocalisations: *Alcedo atthis, Alle alle, Cepphus grille, Botaurus stellaris, Burhinus oedicnemus, Caprimulgus ruficollis, Charadrius hiaticula, Coturnix coturnix, Cuculus canorus, Glareola pratincola, Grus grus, Merops apiaster, Pluvialis apricaria, Pluvialis squatarola, Podiceps nigricollis, Porzana porzana, Tachybaptus ruficollis, Tetrastes bonasia, Upupa epops,* and *Vanellus vanellus*.

We identified a total of 272 species of birds that contribute to mental well-being through their melodious and therapeutic vocalisations.

### 2.2. Assessing anthropogenic threats to NCP provider species

#### 2.2.1. Conservation status of NCP provider species

We used the red list status of species (EEA Red List of Species, 2020) to assess for each NCP, the percentage of provider species in the different threat categories: Least Concern (LC), Data Deficient (DD), Near Threatened (NT), Vulnerable (VU), Endangered (EN), Critically Endangered (CR). We then sought to understand whether the spatial patterns in threatened provider species are regionally distributed or whether this percentage is consistent across Europe, by quantifying the proportion of threatened provider species among the local pool of provider species in each grid cell, separately for regulating and non-material NCP (Fig. 3B).

#### 2.2.2. Major threats to NCP provider species

To understand how threats faced by species can indirectly threaten the NCP they provide, we used the European red list of species dataset. We analysed how major anthropogenic threats indirectly affect NCP, by combining data on species threats (O’Connor et al. 2024) with the species-NCP dataset. The dataset on threats to terrestrial vertebrate species in Europe was derived from the European Red List (EEA, 2018), which provides textual descriptions of threats faced by species across their range. From this source, binary threat information was extracted for each terrestrial vertebrate species by identifying key character strings corresponding to specific threats. Building on the IUCN Red List threat classification scheme (2022), threats were classified into nine broad categories and 20 subcategories. Analysis focused on the six most widespread threat categories, each affecting at least 200 species: urbanization, direct exploitation, agricultural intensification, pollution, invasive alien species (IAS) and diseases, and climate change.

We combined the dataset on threats faced by species and the dataset on provider species for each NCP, to quantify the proportion of provider species that were vulnerable to each major threat type.

### 2.3. Mapping species-mediated NCP across Europe

#### 2.3.1. Study area and species distributions data

The study area covered the spatial extent of the European Union (EU) with the United Kingdom (EU28_+), Norway, Switzerland, and the Western Balkans (Serbia, Kosovo, North Macedonia, Montenegro, Albania, and Bosnia and Herzegovina), matching the study area of the European land systems database (Dou et al., 2021). The NCP maps include all vertebrate species which are known to occur in the study area (EU28+). For this, we used species distribution model (SDM) outputs for all native and breeding terrestrial vertebrate species in the study area and publicly available on the EBV Data Portal (Si-Moussi and Thuiller et al., 2024). Each vertebrate species was modelled using an ensemble of machine learning algorithms. These algorithms were fitted on curated presence-only data from GBIF at 1-km precision with various sets of pseudo-absences, in function of selected environmental data including climate (CHELSA, Karger et al., 2017, 2020), soil (Soilgrids, Poggio et al., 2021), terrain (Amatulli et al., 2018), hydrography (EU-Hydro) and land use (Sandström et al., 2023) variables. In total, Si-Moussi and Thuiller (2024) used 3 algorithms (Random Forest, XGBoost, Neural Networks), 5 pseudo-absence repetitions and 5 spatial block cross-validation partitions (i.e., 75 models). Ensemble mean predictions were made with models that reached a TSS > 0.4 on the cross-validation set. From the provided ensemble predictions available on the EBV Data Portal, we used the constrained predictions which included a spatial constraint based on extent of occurrence range maps (IUCN for mammals and herptiles and BirdLife for birds), which reflects the realised ranges of species. Further details on the species distribution modelling methodology can be found in the Supporting Information.

#### 2.3.2. Mapping NCP capacity

We then mapped species-mediated NCP across Europe, building on species distribution data. To estimate NCP capacity across space, we built on Ceau_u et al. (2021) and measured provider richness in each grid cell as a proxy for NCP capacity. We also explored the biogeography of threatened provider species: we quantified and mapped the percentage of threatened provider species (across all NCP) in each grid cell (Fig. 3B).

In addition to provider richness, we also mapped functional diversity of the provider species for each NCP. To do so, we created functional groups based on species traits using data from Maiorano et al., 2020 and Thuiller et al., 2015, including body mass, diet categories, nesting habitat, feeding space, and feeding behaviour which represent multiple dimensions of species niche. We computed pairwise Gower dissimilarities using the R package cluster, and used hierarchical clustering (Ward’s_D2) on the resulting distance matrix to partition species into 30 clusters that each represented a functional group. We then mapped functional diversity for each NCP by quantifying the number of unique functional groups represented by the provider species that were present in each grid cell for each NCP. We then evaluated the Pearson’s correlation coefficient between functional diversity and provider species richness for each NCP.

#### 2.3.3. Mapping NCP supply

Building on previous work (Verhagen et al., 2017, O’Connor et al., 2021), we defined the supply for each NCP as the product between capacity and demand, after both have been standardized between 0 and 1. To build the demand layer, we built on a previous study by Burkhardt et al., (2014), where the authors applied an expert-based evaluation matrix that assigned scores from 0 to 5 to indicate the societal demand for ecosystem services in each land use class. We used the European land system classification and map (Sandström et al., 2023), and evaluated the extent to which there can be demand for each NCP in each land use class in a table on a scale from 0 to 1 based on the study from Burkhardt et al., (2014) and a literature review beyond (Table S2). In the case of three NCP (species of cultural importance, game species, EDGE), we assumed the demand to be global (culturally important species are valued for their existence): therefore for these NCP, we assume NCP capacity to be equal to the supply, following previous studies (Verhagen et al., 2017; O’Connor et al., 2021). This table was then used as the basis to map the demand across European land systems (Sandström et al., 2023) and further multiply this with the capacity for each NCP, to obtain the NCP supply maps (Fig. 5).

#### 2.3.4. Relationship between NCP multifunctionality and land use intensity

We analysed the relationship between land use intensity and NCP multifunctionality (defined as the capacity of ecosystems to provide multiple benefits to people), in order to assess whether higher land use intensity was associated with lower NCP capacity across multiple NCP. To quantify NCP multifunctionality, we first standardized provider species richness values for each NCP between 0 and 1, using min-max normalization across all grid cells. We then calculated multifunctionality as the average standardized richness across all NCP, separately for regulating and non-material NCP. This provides a measure of the capacity for multiple NCP in each grid cell.

We then fitted a linear model with NCP multifunctionality as the response variable (separately for cultural and regulating), using land use intensity (low, medium, high), land cover type, and their interaction as predictors. We used low-intensity as the reference level for land use intensity, and forests as the reference level for land cover type. All analyses were conducted in R version 4.4.0 (R Core Team, 2024).

## 3. Results

In total, there were 1,858 links between European terrestrial vertebrate species and the 15 NCP (Fig. 1). Out of the 1,168 terrestrial vertebrates considered, 860 species (452 birds, 195 mammals, 78 amphibians, 135 reptiles) provided at least one NCP. Each vertebrate species typically contributed to a limited number of NCP: most species provided between one and three NCP, with a median of 2 NCP per species and a mean of 1.7, and a maximum of 8 different NCP provided by the Eurasian eagle owl (*Bubo bubo*). Species of carnivores, insectivores, raptors and owls typically provided a higher number of NCP than other groups, while water-associated species and small mammals provided one or two NCP (Fig. S3). In terms of conservation status, about 25% of species per NCP were near-threatened or threatened with extinction. Pollinators, ecosystem engineers, seed dispersers, game species, and song birds tended to be less threatened overall than other groups of provider species (Fig. 2A). Species of cultural importance were particularly threatened: evolutionary heritage (provided by EDGE species) were the most threatened in the European red list assessment, followed by species that are important for nature tourism and wildlife watching, as well as species of cultural importance. Among regulating NCP, more than 25% of species that regulate pine processionary moths, carrion elimination and tick control were assessed as threatened with extinction. Across space and combining all NCP, up to 10% of provider species were threatened with extinction (Fig. 2B). The proportion of provider species that are threatened with extinction was particularly high in the Macaronesian islands, Sardinia, southern Italy, the Iberian peninsula, west Ireland, the Balkans coast, and the Scandinavian mountains.

**Figure 2:**
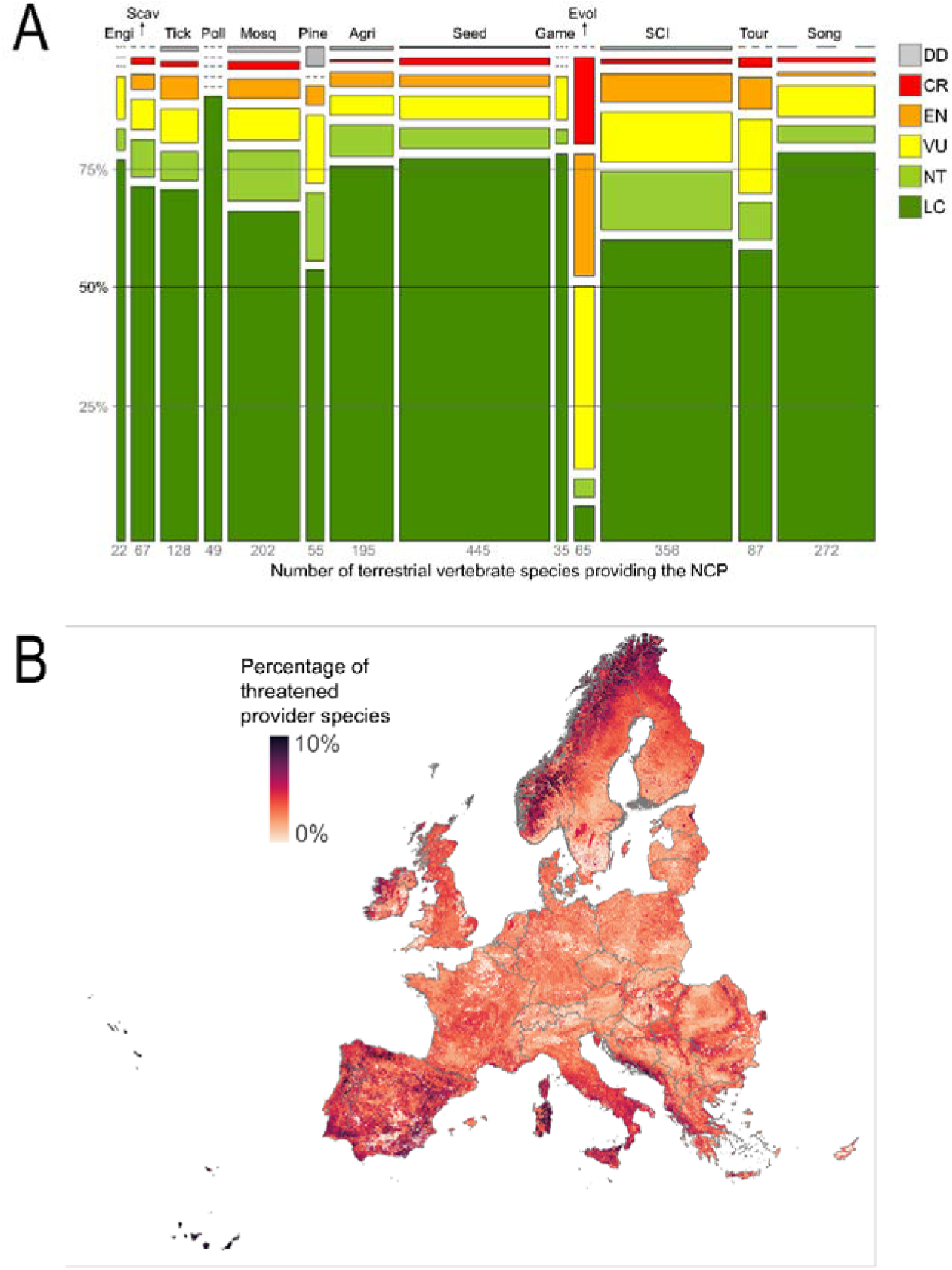
Conservation status of species providing NCP. **(A)** Mosaic plot representing the percentage of provider species that are assessed in different IUCN red list categories, for each NCP, and coloured by threat status (DD: data deficient; CR: critically endangered; EN: endangered; VU: vulnerable; NT: near-threatened; LC: least concern). The width of the columns represent the number of species providing each NCP (shown on the x-axis). We omitted two NCP here: Asian hornet regulation and bark beetle regulation due to very few provider species (2 and 4, respectively). Abbreviations: Engi: ecosystem engineers; Scav: carrion elimination; Tick: tick host control; Pol: pollination; Mosq: mosquito control; Pine: pine moth control; Agri: agricultural pest control; Seed: seed dispersal; Game: game species; Evol: evolutionary heritage; SCI: species of cultural importance; Tour: wildlife tourism; Song: birdsong. **(B)** Percentage of threatened provider species (across all NCP, quantified for each 1km² grid cell across Europe.

Agricultural intensification (especially through pesticide use) was a major threat for 53% of species that provide agricultural pest control (Fig. 3). Direct exploitation was a threat for 74% of game species as well as for 67% of species important for nature tourism and wildlife watching. Direct exploitation also was a major threat to species that can regulate pathogens, impacting 71% of species that eliminate carrion, and 65% of species that regulate tick hosts.

**Figure 3:**
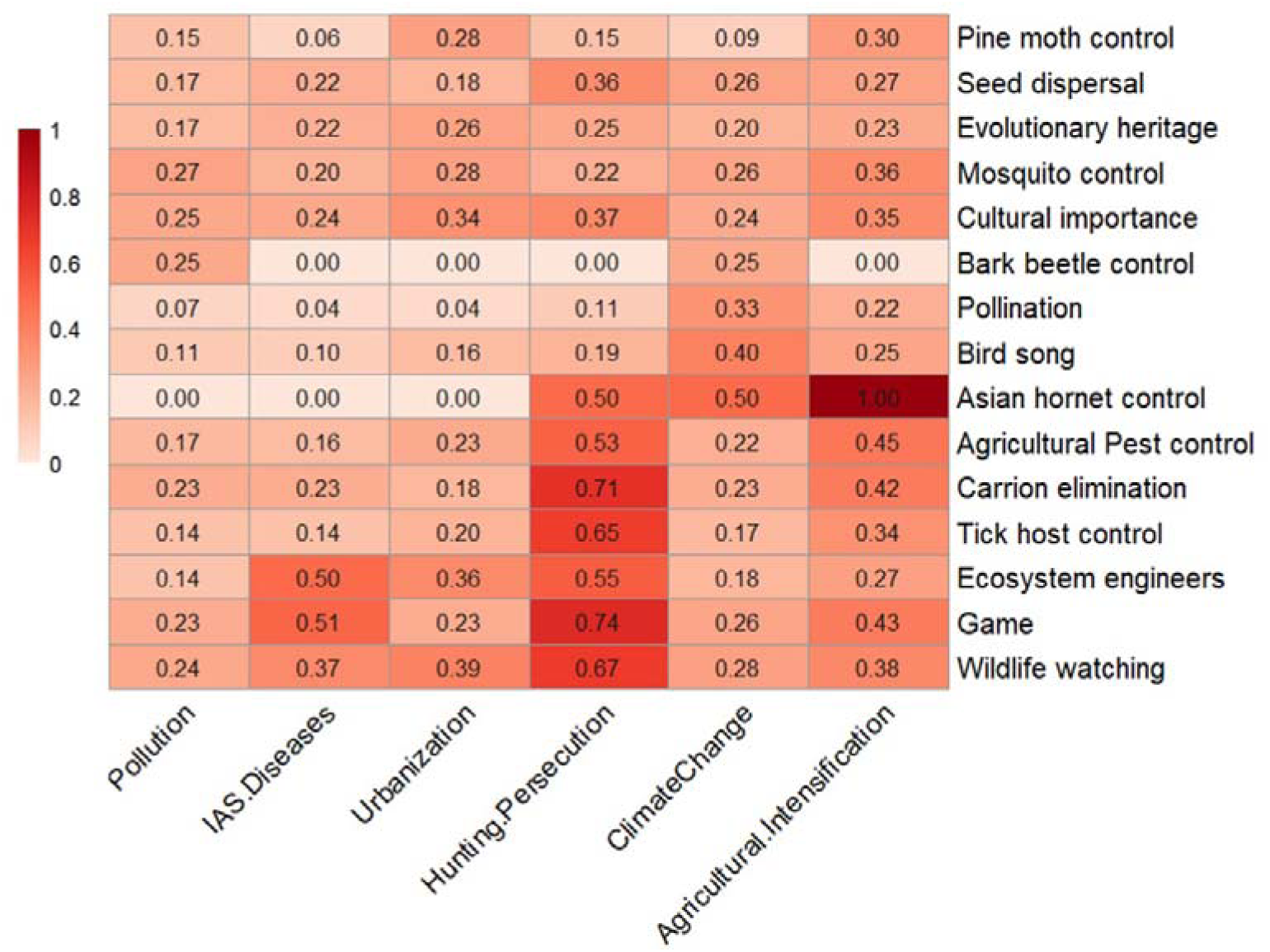
Anthropogenic threats to species-mediated NCP. NCP provided by terrestrial vertebrates are indirectly threatened by human activities. The heatmap shows the proportion of provider species for each NCP (vertical axis) that are affected by six major threat types (horizontal axis). We combined species-mediated NCP with data on species-specific threats (O’Connor et al. 2024). Data on threat types affecting each species was extracted from the European Red list of species.

Local diversity of provider species for both regulating and non-material NCP was significantly lower in high-intensity land use classes, across Europe (Fig. 4). A linear model revealed that land-use intensity and land cover type significantly influenced regulating NCP multifunctionality across Europe (R² = 0.47, p < 0.0001). Regulating NCP multifunctionality was lowest in bare areas, urban areas, high intensity croplands and grasslands, and glaciers and water bodies. Regulating NCP multifunctionality was lower in high intensity classes across land systems, relative to low-intensity forests. Mosaic landscapes (e.g., forest/shrub/grassland and forest/shrub/cropland mosaics) were associated with higher regulating NCP multifunctionality values than low-intensity forests, highlighting the importance of landscape heterogeneity in supporting a diversity of NCP provider species locally. Interaction effects revealed that both medium-intensity arable land and grasslands exhibited higher multifunctionality than expected. Similarly, the model for non-material NCP multifunctionality (R² = 0.42, p < 0.0001) revealed that non-material NCP multifunctionality was lowest in high intensity land systems, in bare areas, and permanent croplands, whereas mosaic systems showed higher non-material NCP multifunctionality values compared to forests, as with regulating NCP. Interaction terms indicated that the negative effects of intensity on non-material NCP multifunctionality were most pronounced in urban areas, while medium-intensity arable land and grasslands showed elevated cultural multifunctionality relative to their low-and high-intensity counterparts.

**Figure 4:**
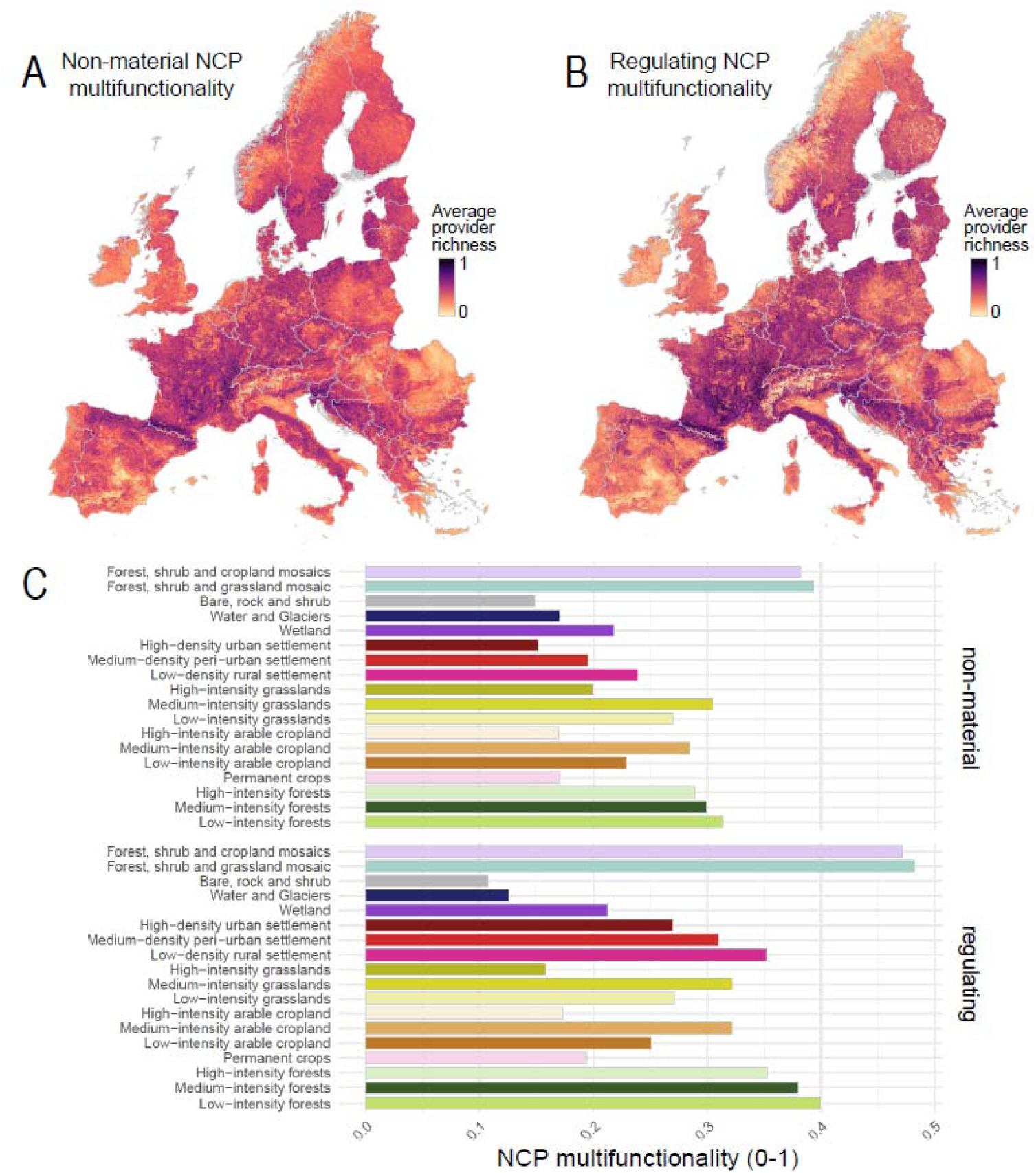
Average provider richness for regulating and non-material NCP across Europe. The maps show the average value of provider species richness of all non-material (left) and regulating (right) NCP in each grid cell, from low (light yellow) to high (dark purple) richness, across Europe (1km²). Species provider richness was normalised between 0 and 1 for each NCP and then averaged across all non-material and regulating NCP, respectively. Species provider richness is a proxy for NCP capacity. The bar plots below represent the average NCP capacity value for all non-material NCP (left) and regulating NCP (right) in each of the European land use intensity classes (Sandström et al., 2023).

Across the 15 NCP, Pearson correlation coefficients between species richness and functional diversity of provider species ranged from r = 0.69 to 0.92. The lowest correlation (0.69) was for pine moth control, while the strongest (0.92) was for nature tourism. These strong to very strong positive relationships indicate that species richness is a reliable proxy for functional diversity across NCP, and spatial patterns of functional diversity tended to resemble those of provider species richness overall (Figs. S6-8).

NCP supply varied substantially between the fifteen NCP categories and across different European regions, reflecting the biogeographic distributions of their provider species as well as spatial variations in societal demand across European land systems (Fig. 5). The NCP supply provided by ecosystem engineers, bark beetle predators, seed dispersers, pine moth predators, wildlife watching tended to be highest in the forested and mountainous regions of Europe. This contrasted with spatial patterns of the NCP supply of agricultural pest control, tick host control, bird song, pollinators, game species, invasive hornet control, comparatively low in these regions, and which tended to be higher in the lowlands. A couple of NCP were highly specific to certain regions of Europe: evolutionary heritage (due to the narrow range size of EDGE species); and mosquito control (due to the high demand in settlements). Overall, this suggested that, across the fifteen NCP considered, mountainous regions, low-intensity forests, and mosaic landscapes act as hotspots of NCP provided by terrestrial vertebrates.

**Figure 5:**
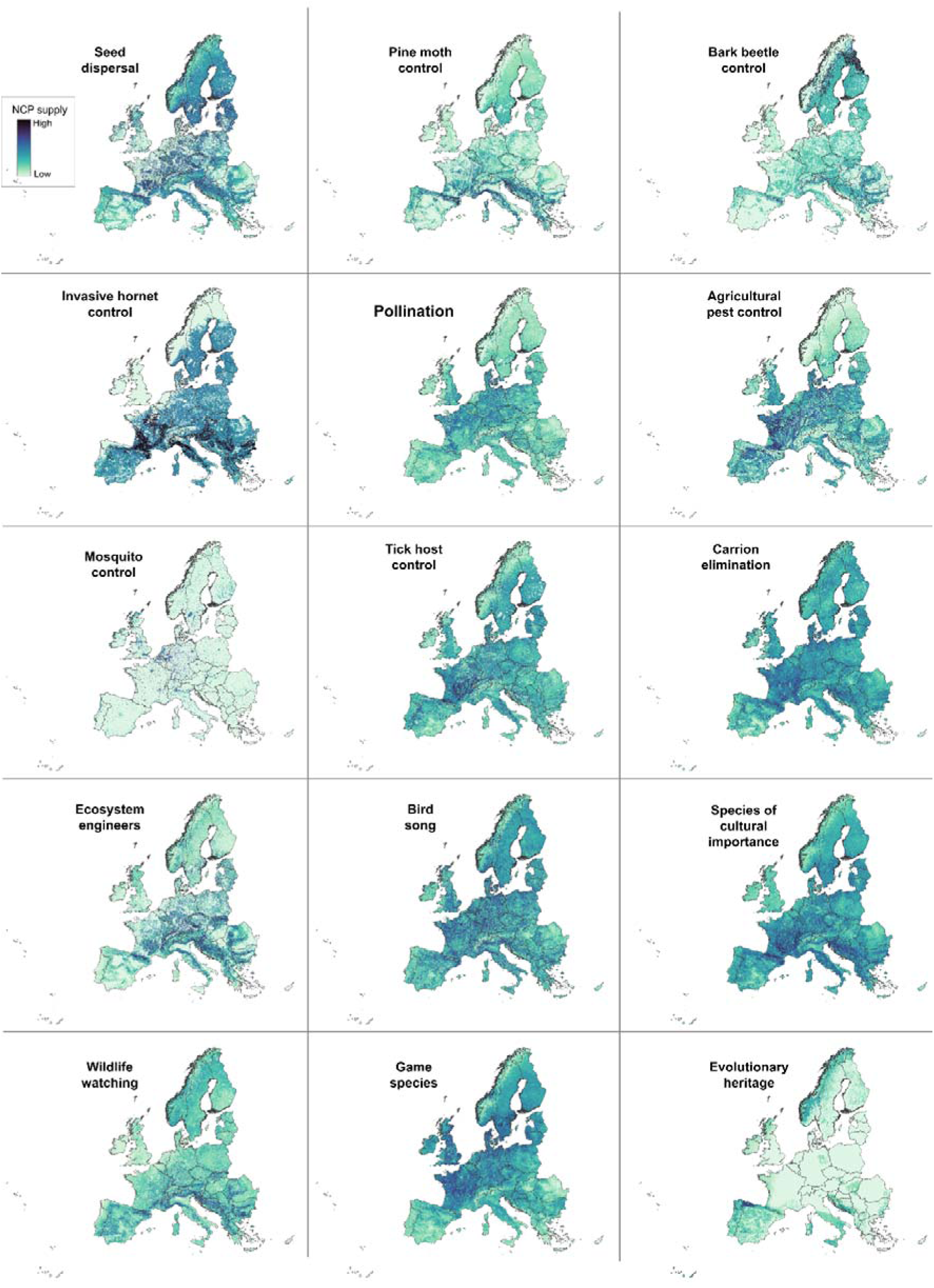
Maps of vertebrate species-mediated NCP supply across Europe (1km²). Supply is measured by capacity demanded (*sensu* Verhagen et al., 2017), as the product of capacity and demand for each NCP.

## 4. Discussion

Here, we synthesised NCP provided by European terrestrial vertebrates, identifying 910 regulating NCP and 5 non-material NCP, and we mapped NCP capacity and supply across Europe. We found that land cover composition and intensity jointly shape the capacity of ecosystems to support a diversity of vertebrate-mediated NCP. The diversity of regulating and non-material NCP provided by terrestrial vertebrates were lower in high intensity land use classes across all land cover classes. Most species that are culturally valued are absent from urban areas, and species that provide regulating NCP are mostly absent from high intensity agricultural landscapes. This is consistent with recent findings that food webs complexity and diversity decreases in high intensity land uses (Botella et al., 2024). We also found an elevated NCP multifunctionality in mosaic land systems (forest/shrub/grassland and forest/shrub/cropland), highlighting the positive role of habitat heterogeneity in supporting a diverse range of species and associated ecological functions and NCP. Interestingly, we found that NCP multifunctionality was higher than expected in medium-intensity croplands and grasslands, which may be due to higher fertiliser inputs than in low-intensity systems and lower disturbance than in high-intensity systems. These findings align with the intermediate disturbance hypothesis (Connell, 1978), and are consistent with evidence that multifunctionality often peaks at intermediate land-use intensities where productivity and habitat heterogeneity can sustain NCP (Lavorel et al., 2011). Furthermore, most species that provide agricultural pest control are strongly threatened by agricultural intensification and pesticide use, which in turn is likely to negatively affect human health and the economy (Frank 2024). This is in line with previous studies showing that organic agriculture promotes pest control (Crowder et al. 2010). Threat mitigation and de-intensification of land use could allow many species to recover, and species-mediated NCP supply to increase, with knock-on socio-economic and health benefits (Frank 2024). This is especially critical in the case of agricultural intensification and direct exploitation which strongly jeopardise the species and ecosystems’ ability to provide many NCP. Our findings thus support the need to restrict or ban direct exploitation of species and to promote regenerative farming practices. Regenerative, organic, and other agroecological systems, such as diversified crop rotations, reduced pesticide use, agroforestry, and the restoration of semi-natural habitats, can enhance habitat heterogeneity, improve soil ecosystem functioning, and increase biodiversity and associated NCP such as pest control and pollination (Torralba et al., 2016; Dainese et al., 2019). Studies have suggested that diversified farming practices in organic agriculture have smaller yield gaps with conventional agriculture than non-diversified systems (Ponisio et al., 2015). Importantly, the EU Nature Restoration Law calls for scaling up such regenerative and agroecological approaches to restore degraded ecosystems, including in and around agricultural landscapes, thereby securing long-term NCP supply.

While we aimed to be as comprehensive as possible in identifying the relationships between vertebrate species and NCP in Europe, our analysis was necessarily constrained by data availability. The species-NCP relationships described in this study are derived from current knowledge contained in published literature and from available datasets on the traits, diets and trophic interactions of these species, which may not be exhaustive. As such, it is likely that many more species contribute to NCP in ways that are not captured here, for example through ecological interactions (Keyes et al., 2021) or functional traits that remain under-documented (Hortal et al., 2015). A key limitation of relying on trophic information is that it assumes spatial and temporal overlap between predators and their prey, which may not always occur in cases of partially overlapping habitats or dietary plasticity of predators; moreover, local NCP supply may be disproportionately influenced by the biomass or abundance of a few key provider species. Finally, complex biotic interaction networks may support key NCP-providing species indirectly, further expanding the set of species involved in sustaining NCP, known as supporting species (Keyes et al., 2021). There is a need for continued research to improve our understanding of the diversity of species underpinning NCP.

In this study, we assumed that provider species richness is a proxy of NCP capacity. Provider species richness is important for effective and resilient provision of the service, particularly for the regulation of zoonoses and pests, and for the pollination of diverse crops. Species richness plays a vital role in long-term NCP supply by increasing redundancy and resilience to fluctuations in the presence or abundance of certain species (Biggs et al. 2020). Predator species richness, for example, is crucial in regulating zoonotic disease reservoirs, such as rodent-borne or mosquito-borne diseases. Optimal disease control is achieved when multiple predator species can simultaneously stabilise local NCP capacity by controlling reservoir populations (Ostfeld & Holt, 2004). However, our approach has certain limitations. First, it should be noted that some species may be more efficient than others (Ceausu et al., 2021) due to a combination of their intrinsic traits (such as body mass, diet specialisation, and preferences) and local density, biomass or resource availability. Provider richness is strongly linked to resilience of the NCP but does not always translate into a higher quantity: the abundance of a single provider species can determine the quantity of NCP provided locally (*e.g.* pollination of crops by *Apis mellifera*). Modelling energy fluxes is a promising tool for estimating the functional effectiveness of species in providing NCP, in the case of regulating NCP that are underpinned by a trophic interaction (Antunes et al. 2024). However, abundance and biomass data are not readily available for all species across Europe, thus we could not account for the relative efficiency of species to provide NCP. Second, our assumption that NCP capacity increases linearly with the number of species providing it may lead to an overestimation, as the relationship may be non-linear (Cardinale et al. 2012). There could be thresholds where a certain level of biodiversity is required to provide a significant amount of NCP; or conversely, there could be situations where competition between functionally similar provider species may decrease NCP supply.

Knowledge of species-mediated NCP can help to predict the vulnerability of NCP to anthropogenic pressures or future scenarios of climate and land use change, and to prioritise species or areas for conservation planning. There are several ways in which such a dataset on species-mediated NCP can be used in conservation planning. NCP supply maps can be used as input features alongside other biodiversity layers (species, habitats) in the same way as other ecosystem service layers such as carbon or water (Jung et al., 2021; O’Connor et al., 2021). Another option is to explicitly incorporate service provision into species-level conservation planning by assigning higher priority (via upweighting) to provider species, thereby aligning biodiversity targets with ecosystem service delivery. Building on the framework developed by Keyes et al. (2021), we could identify keystone species that are important for the robustness of food webs and the supply of NCP across Europe, in particular in future scenarios of climate change and land use changes. This database can thus be useful for identifying priority species that are functionally important and irreplaceable NCP providers, as well as priority areas for conservation and examine the synergies and trade-offs between spatial conservation for species and NCP (Xiao et al. 2018). In the context of restoration planning, areas with high societal demand for an NCP but low NCP provider richness (or abundance), could be prioritised for restoring the NCP provider species, in order to increase NCP value in the areas that matter most. If spatial data on threat occurrence were available, we could map threats to NCP based on the vulnerability of provider species (O’Connor et al. 2024). Better understanding which species provide NCP together with the threats they face can help identify conservation priorities that benefit both nature and people.

