## Supporting Information for "Threats to Nature’s contributions to people provided by terrestrial vertebrates across Europe"

**Extended methods**

**Selection of relevant NCP**

To refine our initial list of species-mediated NCP, we applied the following exclusion criteria. First, we excluded services for which there is insufficient evidence on their prevalence or importance in Europe: wildfire control by large ungulates is a recognized service in African savannahs (Holdo et al., 2009), but its relevance in European ecosystems remains unclear; the aesthetic value of species lacks a standardized metric for European vertebrates; and economic benefits, such as job creation and revenue generation, cannot be easily attributed to specific species. Second, we excluded NCP that were too specific to certain contexts to ensure relevance at the continental scale: frog consumption is culturally important in France (Hocking & Babbitt 2014) but is not broadly relevant across Europe. Third, we excluded services that are frequently accompanied by negative impacts or trade-offs, where the same species or behavior may also be perceived negatively by humans: herbivores that can control weeds may also consume crops and plant seeds in a non-selective manner (Sarabi 2019; Schumacher et al. 2020; Tschumi et al. 2018).

**Detailed methods – Vertebrates SDMs**

**Study extent and target taxa**

Species distributions were modelled at 1 km² resolution for terrestrial vertebrates across Europe. The calibration domain extended beyond the EEA39 reporting region (EU27, the United Kingdom, the four EFTA countries—Iceland, Liechtenstein, Norway, Switzerland—and Turkey) to include adjacent areas in Russia, North Africa, and the Middle East thereby reducing niche truncation (Thuiller et al., 2004; Barbet-Massin et al., 2010).

The initial vertebrate pool comprised 1,259 species across Amphibia, Aves, Mammalia, and Reptilia from the TetraEU database (Maiorano et al. 2020). This list covers most species reported under the EU Habitats (Article 17) and Birds (Article 12) Directives, as well as many species of conservation concern on the IUCN Red List. After data curation, 1,210 species were retained for modelling.

**Environmental covariates**

Environmental covariates were assembled at 1 km² resolution from publicly available datasets. Climatic variables (temperature, precipitation, snow cover trends) were obtained from CHELSA Karger et al. (2017, 2020). Topographic variables (slope, landforms, coastal proximity) were derived from EarthEnv (Amatulli et al 2018). Soil physical and chemical properties (pH, soil organic carbon, texture) were taken from SoilGrids Poggio et al. (2021). Hydrographic layers describing the extent and salinity of water bodies and wetlands were compiled from various land cover products; CORINE Land Cover 2018, ESA CCI Land Cover and the Global Lakes and Wetlands Database (GLWD) Lehner et al. (2004, 2025), while river density was computed using the EU-Hydro (EEA 2019) and HydroSHEDS Lehner et al. (2008) watercourses polygons. Land systems and land-use intensity were obtained from Sandström et al. (2023). To minimise collinearity, only variables with pairwise correlation < 0.70 were retained.

**Occurrence data and pseudo-absences**

Species observations were obtained from GBIF (1980–2023; location uncertainty < 1 km) between 4–9 January 2023^[1]^. Records were cleaned with **CoordinateCleaner** (Zizka et al., 2019) to remove duplicates, records with missing or imprecise coordinates, and common artefacts such as country centroids or institutional localities. Non-avian vertebrates were complemented with IUCN ranges, and birds with breeding ranges from BirdLife International. To account for potential range changes since the last expert assessment, ranges were rasterised and buffered to include 90% of outlier GBIF records within each class. Species with a high proportion of records outside buffered ranges were manually reviewed. Sampling effort maps were generated from observation density per pixel across predefined target groups: Amphibians, Birds, Small Mammals, Large Mammals, Chiroptera and Reptilia.

Pseudo-absences were generated following Barbet-Massin et al. (2012). Three categories were defined: (1) certain presences within buffered ranges, (2) certain absences sampled outside buffered ranges, and (3) uncertain absences sampled within buffered ranges, weighted by target group sampling effort. For 328 species lacking GBIF records but with available expert maps, datasets were generated by sampling certain presence pixels within IUCN ranges proportional to range size and certain absence pixels outside the range. Conversely, for 52 species with GBIF records but no expert range datasets included all observed presences and pseudo-absences. Species with not enough GBIF records (>20 records) and no IUCN ranges were modeled using habitat directives data (n=2) and dropped otherwise (n=49). The procedure was repeated five times per species. Adjustments were made for data-deficient species.

**Species distribution models**

Species distribution models were fitted using Random Forest (RF) Breiman et al. (2001), extreme gradient boosting (XGBoost) Chen et al. (2015), and multilayer perceptron neural networks (MLP) Lecun et al. (2015). Hyperparameters were tuned with GridSearch in scikit-learn Pedregosa et al., (2011). For RF, tree depth and feature subsampling were tuned; for XGBoost, regularisation was additionally optimised; for MLP, architectures of varying depth and width were tested. Class weights were applied to correct for imbalance between presence and absence particularly for rare species.

Presence–pseudoabsence data were partitioned into five spatial blocks (Roberts et al., 2017). Models were trained on four blocks and tested on the fifth. Only models with cross-validated True Skill Statistic (TSS) > 0.4 were retained. Ensembles combined models across algorithms, folds and pseudo-absence replicates.

**Ensemble forecasting**

After calibration and validation, SDMs were projected to maps of environmental suitability ranging from 0 (unsuitable) to 1 (highly suitable). Ensembles were built from all models that met the performance threshold (TSS ≥ 0.4) and combined into a single output: the mean suitability across retained models.

**Post-hoc spatial constraints**

Suitability maps were constrained with expert ranges using the **Estar framework** (Hoareau et al., in prep). Post-hoc spatial constraints were applied by clipping suitability maps to IUCN/BirdLife extent-of-occurrence ranges, smoothed with an exponential decay kernel from range boundaries to buffer limits.

**Threshold optimisation procedure**

For each species, thresholds were derived using the Continuous Boyce Index calibration curve (Hirzel et al., 2006). The calibration curve compares the frequency of presences in predicted suitability bins with the frequency expected under random sampling. Ratios > 1 indicate that presences occur more often than expected, ratios ≈ 1 indicate frequencies close to random expectation, and ratios < 1 indicate fewer presences than expected. From this curve, two thresholds (th_low and th_high) were identified to delimit unsuitable, uncertain and suitable probability ranges. In the constrained maps, only pixels with values above th_high were retained, while the remaining were set to zero.

**Supplementary Figures**

**Figure S1: Overview of regulating NCP provided by terrestrial vertebrates in Europe.**

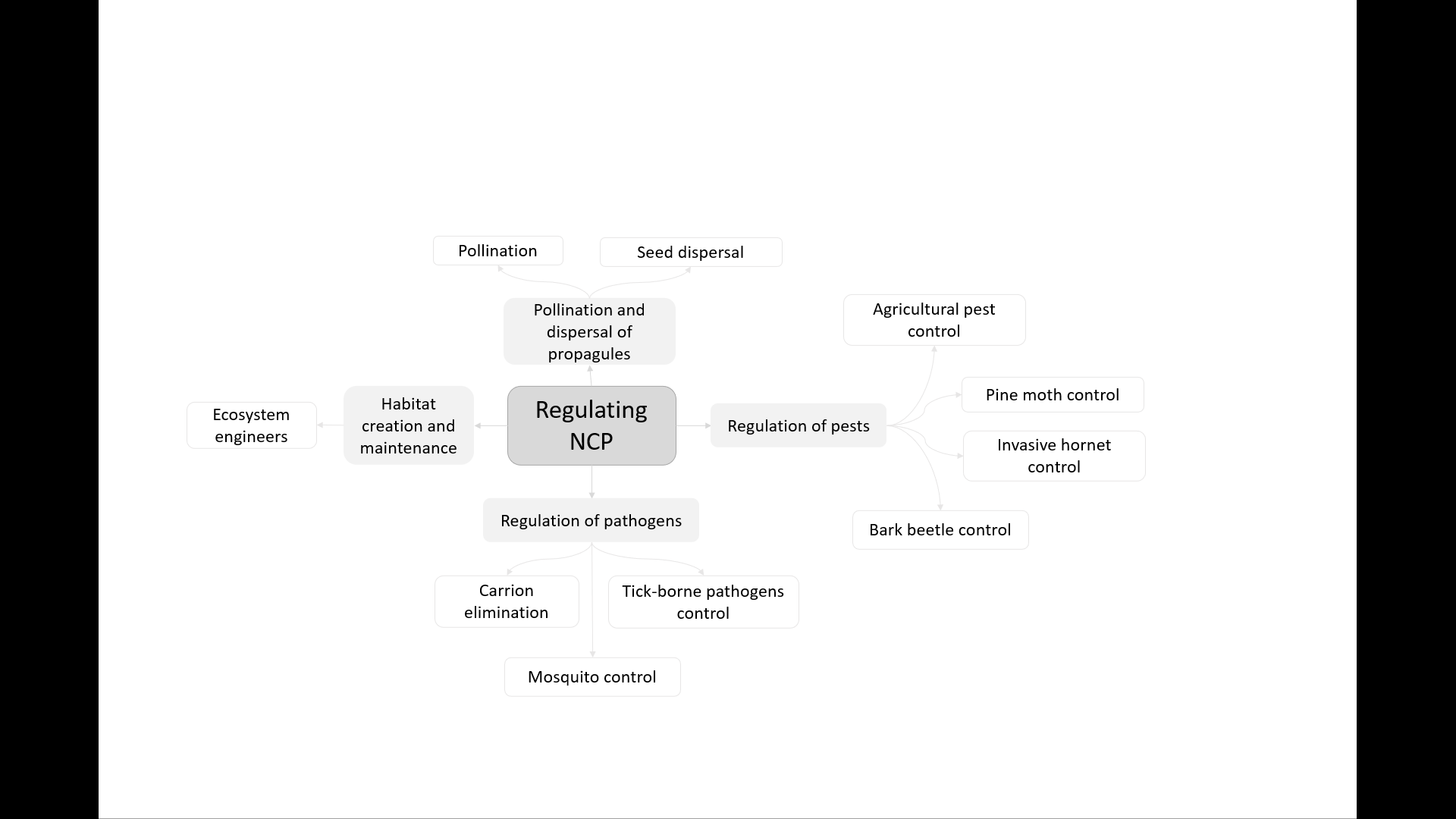

**Figure S2: Overview of cultural NCP provided by terrestrial vertebrates in Europe.**

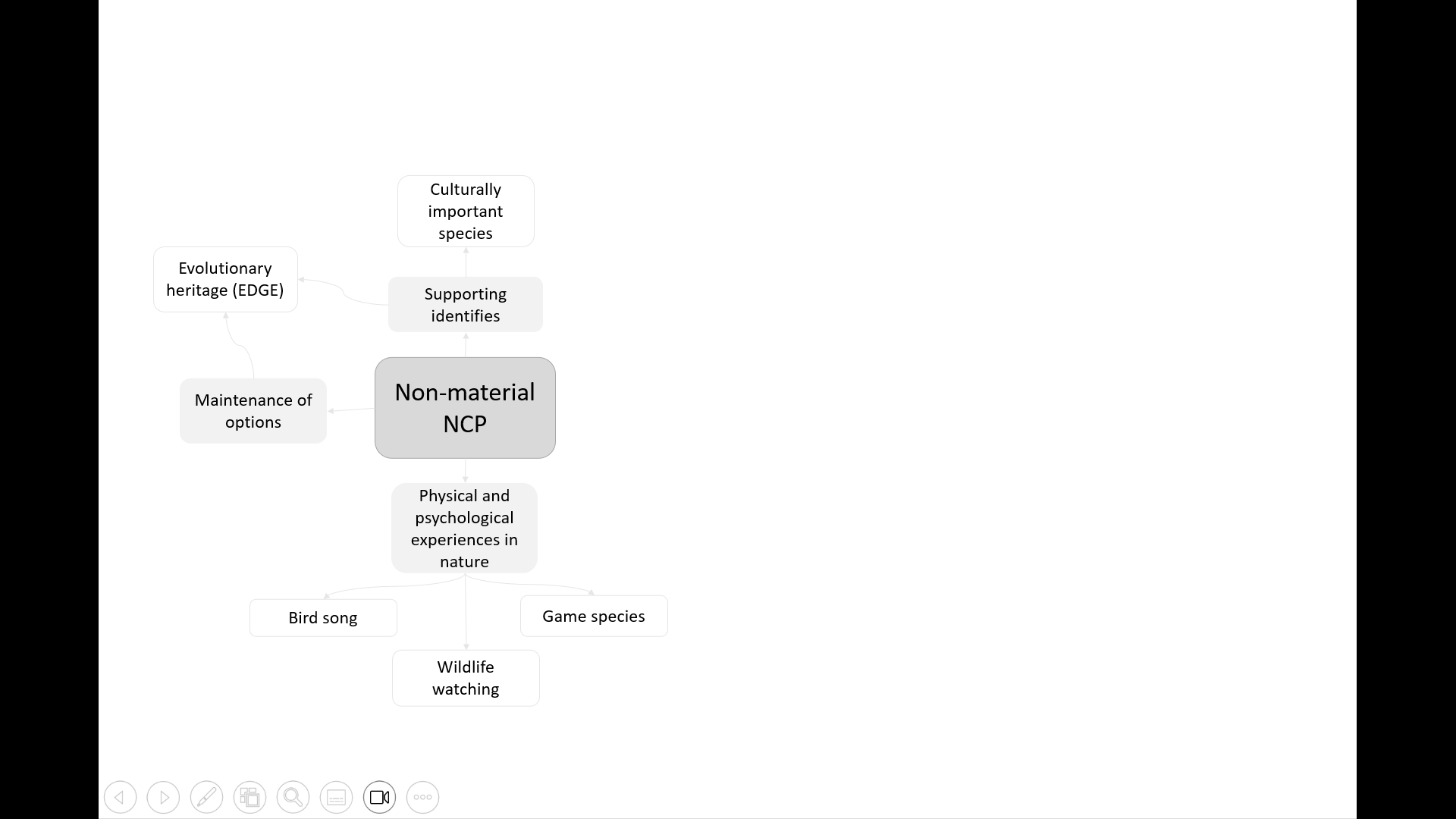

**Figure S3:** **Number of different NCP provided per species across functional groups.**

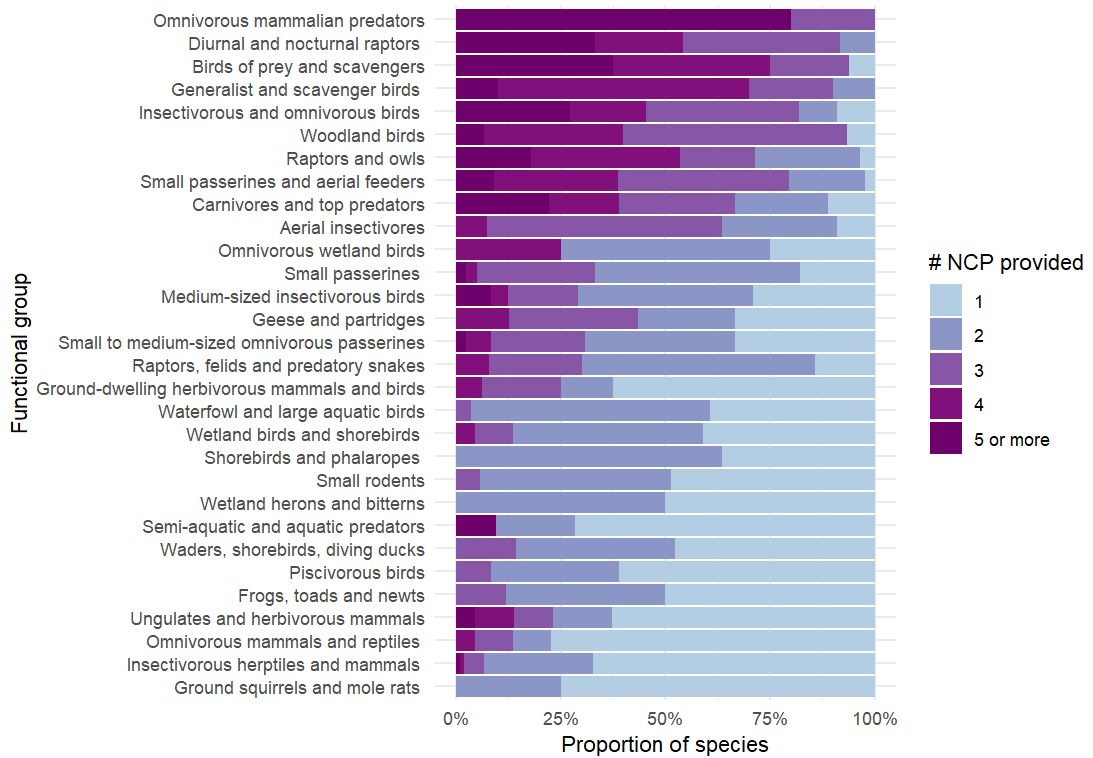

**Figure S4: Identifying top species of cultural importance by extracting the inflection point in the number of Wikipedia page visitation for each species.**

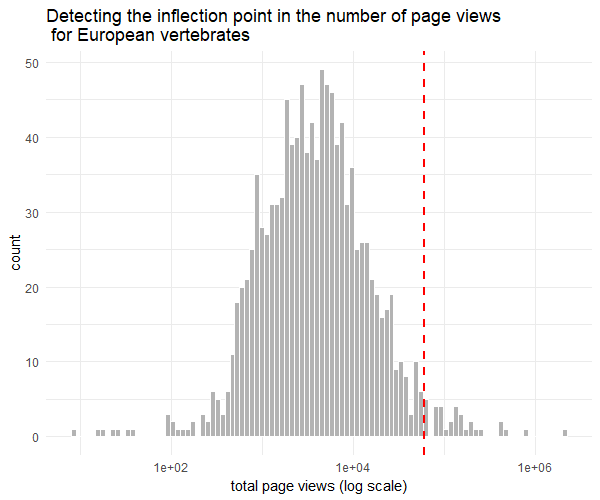

**Figure S5: Identifying top species for wildlife watching by extracting the inflection point in observation density for European vertebrates**

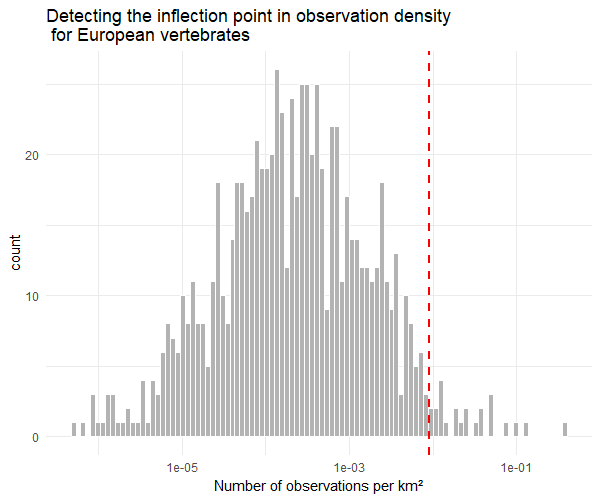

**Figure S6: Maps of species richness and functional diversity for regulating NCP in agricultural and forested systems**

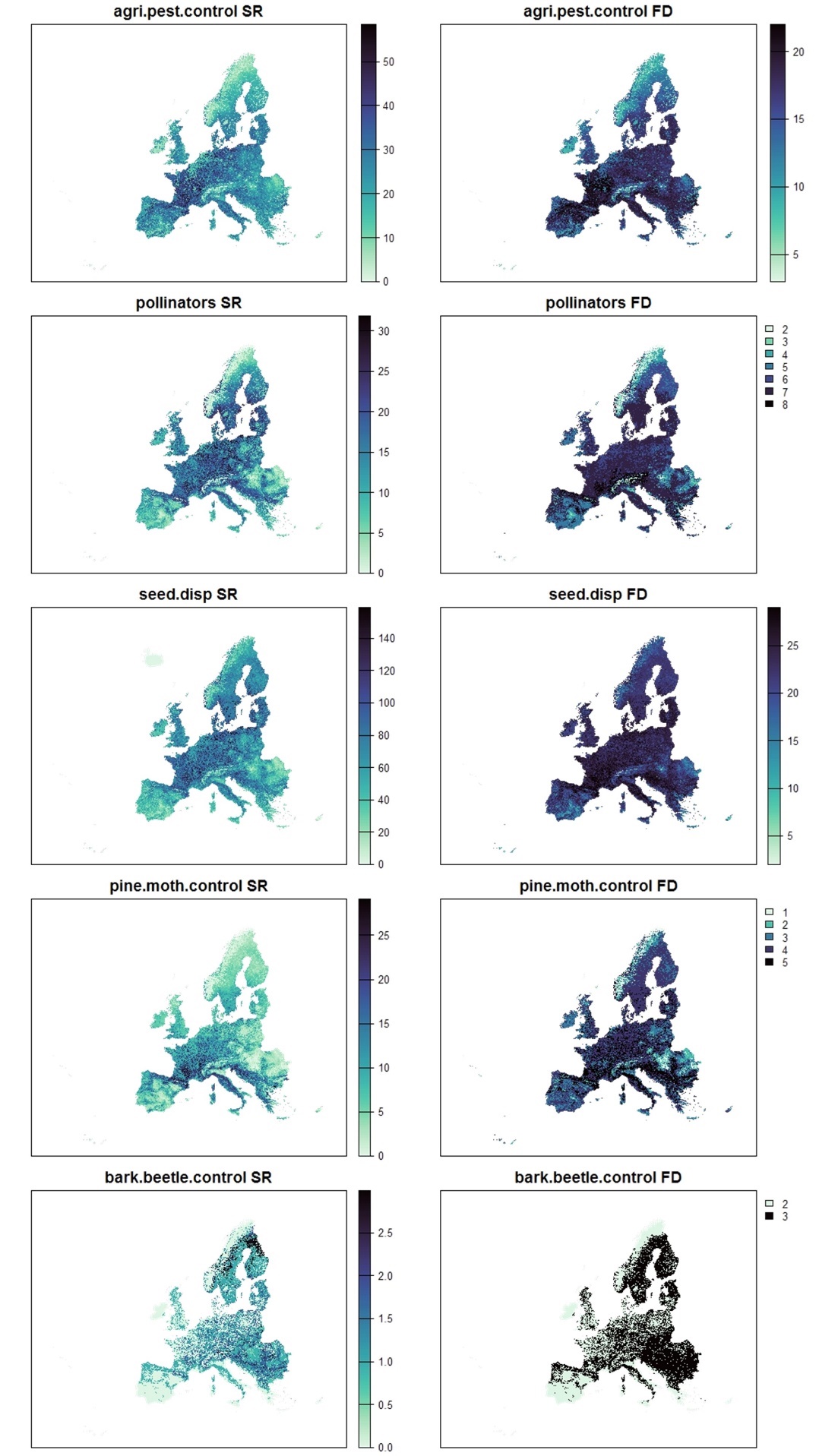

**Figure S7: Maps of species richness and functional diversity for regulating NCP related to habitat maintenance and regulation of diseases and pathogens. Regulation of invasive hornet was omitted from the functional diversity analysis since only two species provide this NCP.**

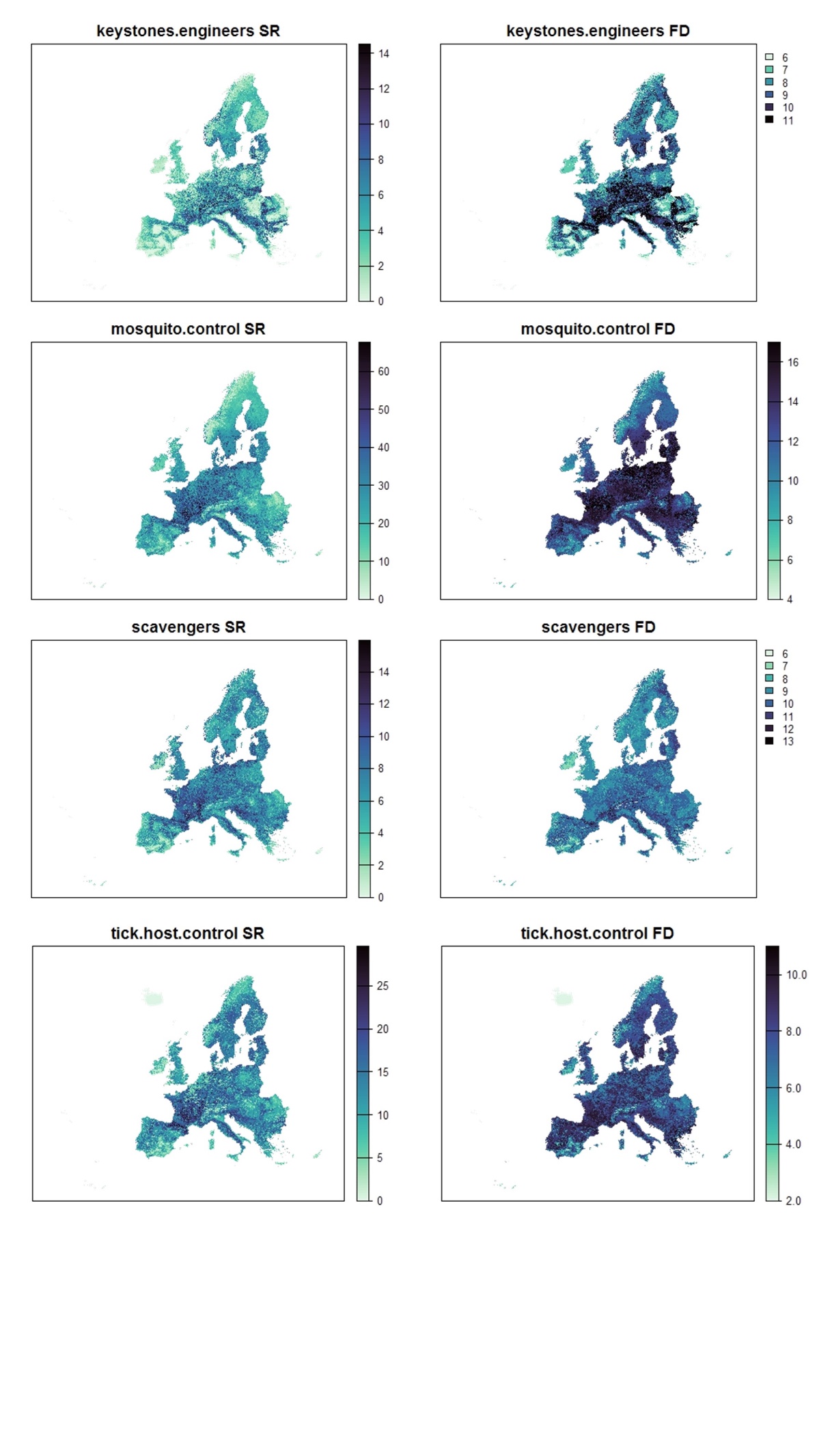

**Figure S8: Maps of species richness and functional diversity for non-material NCP**

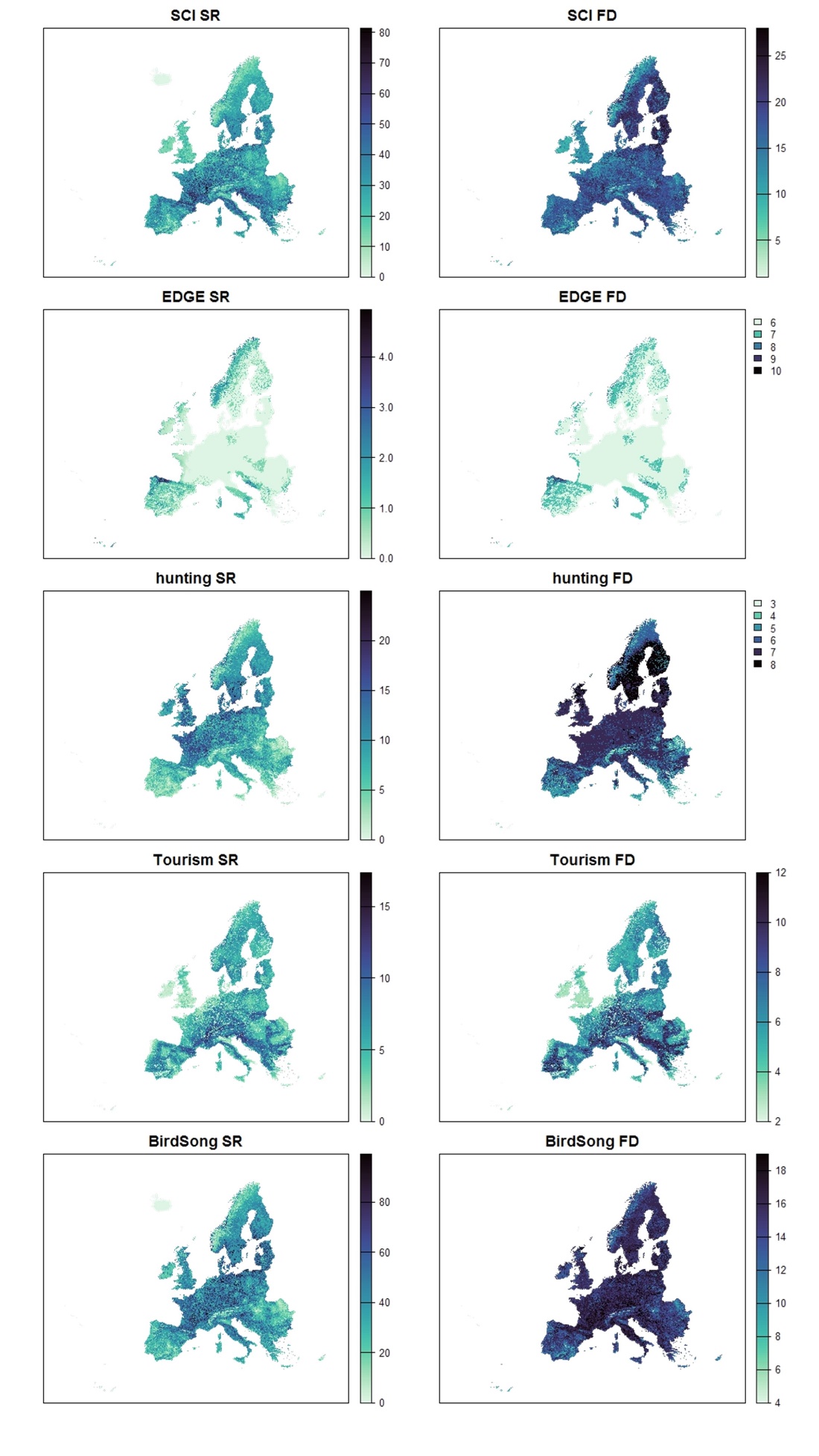

**Table S1:** List of 68 species that feature frequently and prominently on European wildlife tourism websites. These are flagship and/or charismatic species that motivate wildlife watching tourism.

| **Species** | **Common name** | **Wildlife tourism website** | **Country** |
| --- | --- | --- | --- |
| ***Aegolius funereus*** | Tengmalm's owl | https://www.wildlifeworldwide.com/locations/saaremaa-island ; https://www.wildlifeworldwide.com/group-tours/spring-birding-in-slovakia ; https://swedenfishingandbirding.net/ | Estonia, Slovakia, Sweden |
| ***Aegypius monachus*** | Cinereous vulture | neophron.com ; https://wildsideholidays.co.uk/ | Spain, Bulgaria |
| ***Alca torda*** | Razorbill | https://www.wildlifeworldwide.com/group-tours/wildlife-of-lewis-harris | Scotland |
| ***Alcedo atthis*** | Common kingfisher | https://www.wildlifeworldwide.com/group-tours/romanias-wildlife-highlights | Romania |
| ***Alces alces*** | Moose | https://www.wildsweden.com/about/the-wild-animals; https://www.responsibletravel.com/holidays/europe/travel-guide/top-10-wildlife-holidays-in-europe ; https://www.nationalgeographic.com/travel/article/where-to-see-wildlife-in-europe-this-summer ; https://www.wildlifeworldwide.com/discover/europe | Sweden, Poland; Finland |
| ***Alopex lagopus*** | Arctic fox | https://www.wildsweden.com/about/the-wild-animals ; https://www.wildlifeworldwide.com/discover/norway | Sweden, Norway |
| ***Aquila adalberti*** | Spanish imperial eagle | https://wildsideholidays.co.uk/cinereous-vulture-reintroduction-program-takes-flight-in-spains-iberian-highlands/ ; https://www.wildlifeworldwide.com/group-tours/iberian-lynx-raptor-photography | Spain |
| ***Aquila chrysaetos*** | Golden eagle | sustainability.spain.info; https://www.wildlifeworldwide.com/group-tours/islay-jura-in-autumn ; https://www.responsibletravel.com/holidays/europe-wildlife/travel-guide/rewilding; https://www.responsibletravel.com/holidays/europe/travel-guide/top-10-wildlife-holidays-in-europe ; https://www.wildlifeworldwide.com/trip-ideas/golden-eagles-in-winter | Spain, Scotland, Romania, Ukraine, Moldova, Sweden |
| ***Aquila fasciata*** | Bonelli's eagle | sustainability.spain.info ; wildlatitudes.com | Spain |
| ***Aquila heliaca*** | Estern imperial eagle | https://www.wildlifeworldwide.com/group-tours/spring-birding-hungary ; https://www.rockjumperbirding.com/tour-info/croatia-slovenia-austria-bird-photography-tour-2026/50229/ | Hungary, Austria |
| ***Athene noctua*** | Little owl | https://www.wildlifeworldwide.com/trip-ideas/birding-in-portugals-algarve-alentejo | Portugal |
| ***Bison bonasus*** | European bison | https://www.responsibletravel.com/holiday/24699/bialowieza-forest-wildlife-tour-in-poland ; https://www.wildlifenomads.com/blog/fauna-animals-in-europe ; https://www.wildlifeworldwide.com/group-tours/polands-winter-wildlife | Poland |
| ***Bubo bubo*** | Eurasian eagle owl | https://www.birdinginportugal.com/cuckoos-owls-nightjars/eurasian-eagle-owl-bubo-bubo ; https://www.wildlifenomads.com/blog/fauna-animals-in-europe ; https://www.wildlifeworldwide.com/group-tours/polands-winter-wildlife | Portugal , Poland |
| ***Calidris pugnax*** | Ruff | https://www.apex-expeditions.com/expeditions/finland-romania-wildlife-tour/ ; https://www.wildlifeworldwide.com/locations/biebrza-marshes | Finland ; Poland |
| ***Canis aureus*** | Golden jackal | https://www.wildlifeworldwide.com/group-tours/romanias-wildlife-highlights; https://www.birdwatchingholidays.com/tours/golden jackal european souslik wildlife photography?tab=prices | Romania, Bulgaria |
| ***Canis lupus*** | Wolf | https://theeuropeannaturetrust.com/travel/; https://www.responsibletravel.com/holidays/europe/travel-guide/top-10-wildlife-holidays-in-europe ; https://www.nationalgeographic.com/travel/article/where-to-see-wildlife-in-europe-this-summer; https://www.wildlifenomads.com/blog/fauna-animals-in-europe | Sweden, Poland, France; Italy ; Romania |
| ***Capra ibex*** | Alpine ibex | https://www.myswitzerland.com/en-us/experiences/summer-autumn/excursions/wildlife-watching/wildlife-watching/alpine-ibex/; https://www.nationalgeographic.com/travel/article/where-to-see-wildlife-in-europe-this-summer ; https://www.wildlifeworldwide.com/discover/europe | Switzerland; France; Austria |
| ***Capra pyrenaica*** | Spanish ibex | https://wildsideholidays.co.uk/spanish-ibex-capra-pyrenaica-hispanica-cabra-montes/; https://www.responsibletravel.com/holidays/europe/travel-guide/top-10-wildlife-holidays-in-europe | Spain |
| ***Castor fiber*** | Eurasian beaver | https://www.wildsweden.com/about/the-wild-animals/ ; https://wildsideholidays.co.uk/ ; https://www.austria.info/en-gb/activities/animal-experiences/#wildlife-watching ; https://www.responsibletravel.com/holidays/europe/travel-guide/top-10-wildlife-holidays-in-europe ; https://www.nationalgeographic.com/travel/article/where-to-see-wildlife-in-europe-this-summer ; https://www.wildlifeworldwide.com/group-tours/polands-winter-wildlife | Sweden, Spain, Austria, Poland |
| ***Cervus elaphus*** | Red deer | https://rhodopemountains.eu/tours/mammals-birds/; https://www.wildlifeworldwide.com/group-tours/polands-winter-wildlife ; wildlifenomads.com/blog/fauna-animals-in-europe; https://www.wildlifeworldwide.com/group-tours/islay-jura-in-autumn | Bulgaria, Poland, Scotland |
| ***Ciconia ciconia*** | White stork | https://danubedelta.org/en/wildlife/white-stork/ ; https://www.wildlifeworldwide.com/trip-ideas/birding-in-portugals-algarve-alentejo | Romania, Portugal |
| ***Ciconia nigra*** | Black stork | iberianwildlife.com ; sustainability.spain.info; https://www.wildlifeworldwide.com/group-tours/spring-birding-hungary ; https://www.wildlifeworldwide.com/group-tours/spring-birding-in-slovakia | Spain, Hungary ; Slovakia |
| ***Coracias garrulus*** | European roller | https://www.wildlifeworldwide.com/group-tours/spring-birding-hungary ; https://www.wildlifeworldwide.com/group-tours/the-unknown-adriatic/gallery | Hungary , Montenegro |
| ***Dama dama*** | Fallow deer | https://rhodopemountains.eu/tours/mammals-birds | Bulgaria |
| ***Dryocopos martius*** | Black woodpecker | https://swedenfishingandbirding.net/2020/02/18/black-woodpecker-dryocopus-martius-viewing-from-our-private-forest-hide/ ; https://www.rockjumperbirding.com/tour-info/croatia-slovenia-austria-bird-photography-tour-2026/50229/ ; https://www.rockjumperbirding.com/tour-info/bulgaria-greece-spring-in-the-balkans-2026/50180/ | Bulgaria; Sweden ; Austria ; Croatia |
| ***Falco vespertinus*** | Red footed falcon | https://www.wildlifeworldwide.com/group-tours/spring-birding-hungary ; https://www.wildlifeworldwide.com/locations/kiskunsag-national-park | Hungary |
| ***Fratercula arctica*** | Puffin | https://www.responsibletravel.com/holidays/europe/travel-guide/top-10-wildlife-holidays-in-europe ; https://www.nationalgeographic.com/travel/article/where-to-see-wildlife-in-europe-this-summer | Scotland, Iceland |
| ***Gavia arctica*** | Black-throated loon | https://www.wildlifeworldwide.com/group-tours/wildlife-of-lewis-harris | Scotland |
| ***Grus grus*** | Common crane | https://www.wildlifeworldwide.com/group-tours/spring-birding-hungary | Hungary |
| ***Gulo gulo*** | Wolverine | https://www.wildlifenomads.com/blog/fauna-animals-in-europe ; https://www.wildlifeworldwide.com/group-tours/european-festival-of-bears ; https://www.wildlifeworldwide.com/locations/viiksimo ; https://www.apex-expeditions.com/expeditions/finland-romania-wildlife-tour/ | Finland |
| ***Gypaetus barbatus*** | Bearded vulture | https://www.wildlifeworldwide.com/discover/spain ; https://www.responsibletravel.com/holidays/europe/travel-guide/top-10-wildlife-holidays-in-europe ; https://www.nationalgeographic.com/travel/article/where-to-see-wildlife-in-europe-this-summer | Spain, France |
| ***Gyps fulvus*** | Griffon vulture | https://www.knaturewildlife.com/en/daytrips/sardinia-birdwatching-day-trips/ ; naturewatchingineurope.com ; https://www.nathab.com/europe | Spain, Portugal, Italy |
| ***Haliaeetus albicilla*** | White-tailed eagle | https://www.wildlifeworldwide.com/group-tours/northern-lights-lapland-birds ; https://www.wildlifeworldwide.com/group-tours/early-spring-in-the-scottish-highlands https://www.wildlifeworldwide.com/group-tours/polands-winter-wildlife; https://www.rockjumperbirding.com/tour-info/croatia-slovenia-austria-bird-photography-tour-2026/50229/ | Finland, Poland, Scotland, Austria |
| ***Hieraaetus pennatus*** | Booted eagle | https://www.wildlifeworldwide.com/trip-ideas/birding-in-portugals-algarve-alentejo | Portugal |
| ***Lagopus lagopus*** | Willow ptarmigan | [https://www.aladdin.st/bird-watching/sweden/willow ptarmigan.html](https://www.aladdin.st/bird-watching/sweden/willow_ptarmigan.html) | Sweden |
| ***Lagopus muta*** | Rock ptarmigan | https://www.wildlifeworldwide.com/group-tours/early-spring-in-the-scottish-highlands | Scotland |
| ***Lutra lutra*** | Eurasian Otter | https://www.responsibletravel.com/holidays/europe/travel-guide/top-10-wildlife-holidays-in-europe ;https://www.wildlifeworldwide.com/locations/monfraguee-national-park | Poland, Romania |
| ***Lynx lynx*** | Eurasian lynx | https://www.wildsweden.com/about/the-wild-animals/; https://www.responsibletravel.com/holidays/europe/travel-guide/top-10-wildlife-holidays-in-europe ; https://www.wildlifeworldwide.com/group-tours/polands-winter-wildlife | Sweden; Poland; Romania |
| ***Lynx pardinus*** | Iberian lynx | https://theeuropeannaturetrust.com/travel/iberian-lynx-recovery-adventure/ ; https://wildsideholidays.co.uk/mammals-of-spain/ ; https://www.wildlifenomads.com/blog/fauna-animals-in-europe ; https://www.wildlifeworldwide.com/discover/spain | Spain |
| ***Lyrurus tetrix*** | Black grouse | https://www.wildlifeworldwide.com/discover/scotland | Scotland |
| ***Marmota marmota*** | Alpine marmot | https://www.austria.info/en-gb/activities/animal-experiences/#wildlife-watching ; https://www.responsibletravel.com/holidays/europe/travel-guide/top-10-wildlife-holidays-in-europe | Austria, Spain |
| ***Martes martes*** | Pine marten | https://www.wildlifeworldwide.com/discover/netherlands ; https://www.wildlifeworldwide.com/group-tours/polands-winter-wildlife ; https://www.wildlifeworldwide.com/group-tours/early-spring-in-the-scottish-highlands | Netherlands, Poland, Scotland |
| ***Merops apiaster*** | European bee-eater | https://www.wildenatur.com/en/birds/european-bee-eater-merops-apiaster; https://www.wildlifeworldwide.com/trip-ideas/birding-in-portugals-algarve-alentejo ; https://www.wildlifeworldwide.com/group-tours/romanias-wildlife-highlights; https://www.wildlifeworldwide.com/discover/hungary | Germany , Portugal, Hungary, Romania |
| ***Mustela lutreola*** | European mink | https://www.wildlifeworldwide.com/group-tours/romanias-wildlife-highlights | Romania |
| ***Mustela putorius*** | Polecat | https://www.wildlifeworldwide.com/discover/netherlands | Netherlands |
| ***Neophron percnopterus*** | Egyptian vulture | https://www.birdwatchingholidays.com/ ; neophron.com ; https://www.wildlifeworldwide.com/discover/europe | Spain, Bulgaria |
| ***Otis tarda*** | Great bustard | https://www.algarve-birdwatching.com/GreatBustard.html ; https://www.wildlifeworldwide.com/group-tours/spring-birding-hungary ; https://www.wildlifeworldwide.com/discover/spain | Portugal ; Hungary; Spain |
| ***Ovis aries*** | Mouflon | https://wildlifeexperiencespain.com/species/mouflon-sighting-in-spain | Spain |
| ***Pandion haliaetus*** | Osprey | https://www.wildlifeworldwide.com/discover/scotland; https://www.wildlifeworldwide.com/locations/lake-kerkini | Scotland; Greece |
| ***Pelecanus crispus*** | Dalmatian pelican | https://birdinginalbania.com/natural-albania-bird-tour/ ; https://www.wildlifetrails.co.uk/europe-wildlife-holidays/ ; https://www.wildlifeworldwide.com/group-tours/the-unknown-adriatic | Albania, Greece , Montenegro |
| ***Pelecanus onocrotalus*** | Great white pelican | https://sustainability.spain.info/en/top/wildlife-spotting-safari-europe; https://www.wildlifeworldwide.com/group-tours/romanias-wildlife-highlights ; | Spain , Romania |
| ***Phoenicopterus roseus*** | Greater flamingo | https://www.knaturewildlife.com/en/daytrips/sardinia-birdwatching-day-trips/; wildlifenomads.com/blog/fauna-animals-in-europe | Italy , Spain |
| ***Platalea leucorodia*** | Eurasian spoonbill | https://www.wildlifeworldwide.com/group-tours/spring-birding-hungary; https://www.wildlifeworldwide.com/group-tours/romanias-wildlife-highlights | Hungary, Romania |
| ***Polysticta stelleri*** | Steller's eider | https://www.wildlifeworldwide.com/discover/europe | Estonia |
| ***Rangifer tarandus*** | Reindeer | https://www.taigaspirit.com/wildlife-experiences/; https://www.wildlifenomads.com/blog/fauna-animals-in-europe ; https://www.wildlifeworldwide.com/group-tours/northern-lights-lapland-birds | Finland |
| ***Rupicapra rupicapra*** | Chamois | https://www.parconazionale-stelvio.it/en/experiences/guided-nature-experiences/; https://www.nationalgeographic.com/travel/article/where-to-see-wildlife-in-europe-this-summer ; https://www.wildlifenomads.com/blog/fauna-animals-in-europe; https://www.wildlifeworldwide.com/discover/europe ; https://www.wildlifeworldwide.com/group-tours/spring-birding-in-slovakia | Italy , France, Greece; Austria; Slovakia |
| ***Somateria mollissima*** | Common eider | https://www.wildlifeworldwide.com/group-tours/northern-lights-lapland-birds ; https://www.wildlifeworldwide.com/group-tours/islay-jura-in-autumn | Finland, Scotland |
| ***Somateria spectabilis*** | King eider | https://www.wildlifeworldwide.com/group-tours/northern-lights-lapland-birds | Finland |
| ***Strix nebulosa*** | Grey owl | https://www.wildsweden.com/about/the-wild-animals/; https://swedenfishingandbirding.net/ | Sweden |
| ***Strix uralensis*** | Ural owl | https://www.wildlifeworldwide.com/group-tours/spring-birding-hungary ; https://www.wildlifeworldwide.com/group-tours/spring-birding-in-slovakia; https://www.wildlifeworldwide.com/locations/dinaric-alps ; https://www.rockjumperbirding.com/tour-info/croatia-slovenia-austria-bird-photography-tour-2026/50229/ | Hungary ; Slovakia ; Slovenia |
| ***Surnia ulula*** | Northern hawk-owl | https://www.wildsweden.com/about/the-wild-animals/ | Sweden |
| ***Sus scrofa*** | Wild boar | https://www.wildsweden.com/about/the-wild-animals/ ; https://www.wildlifeworldwide.com/group-tours/polands-winter-wildlife ; wildlifenomads.com/blog/fauna-animals-in-europe ; https://www.wildlifeworldwide.com/discover/europe | Sweden, Spain ; Austria, Poland |
| ***Tetrao urogallus*** | Western capercaillie | https://www.taigaspirit.com/wildlife-experiences/ ; https://wildsideholidays.co.uk/tetrao-urogallus-cantabricus-cantabrian-capercaillie-urogallo-cantabrico/ ; https://www.wildlifenomads.com/blog/fauna-animals-in-europe; https://www.wildlifeworldwide.com/group-tours/spring-birding-in-slovakia; https://swedenfishingandbirding.net/ | Sweden, Spain; Slovakia |
| ***Tetrax tetrax*** | Little bustard | https://www.wildlifeworldwide.com/locations/kiskunsag-national-park | Hungary |
| ***Tichodroma muraria*** | Wallcreeper | https://www.wildlifeworldwide.com/group-tours/romanias-wildlife-highlights ; https://www.rockjumperbirding.com/tour-info/bulgaria-greece-spring-in-the-balkans-2026/50180/ | Romania; Bulgaria |
| ***Upupa epops*** | European hoopoe | https://www.wildlifeworldwide.com/trip-ideas/birding-in-portugals-algarve-alentejo ; https://www.rockjumperbirding.com/tour-info/croatia-slovenia-austria-bird-photography-tour-2026/50229/ | Portugal, Austria, Croatia, Slovenia |
| ***Ursus arctos*** | Brown bear | https://www.responsibletravel.com/holidays/europe-wildlife/travel-guide/rewilding; https://www.nationalgeographic.com/travel/article/where-to-see-wildlife-in-europe-this-summer; https://www.responsibletravel.com/holidays/europe/travel-guide/top-10-wildlife-holidays-in-europe; https://www.nationalgeographic.com/travel/article/where-to-see-wildlife-in-europe-this-summer ; https://www.wildlifenomads.com/blog/fauna-animals-in-europe; https://www.wildlifeworldwide.com/discover/slovenia; https://www.apex-expeditions.com/expeditions/spain-wildlife-tours/ | Spain, Sweden; Romania; Slovenia |
| ***Ursus maritimus*** | Polar bear | https://www.nationalgeographic.com/travel/article/where-to-see-wildlife-in-europe-this-summer; https://www.wildlifenomads.com/blog/fauna-animals-in-europe ; https://www.wildlifeworldwide.com/discover/norway | Norway |

**Table S2:** Demand for each NCP in each land use class. A value of 1 (coloured in green) indicates that a land use class is associated with a high demand for the NCP. A value of zero (coloured in red) indicated that there is no demand for the NCP in the land use class. In the case of three cultural NCP (species of cultural importance, game species, EDGE), the demand is global (these species are culturally valued for their existence), or the demand does not vary across land use classes: therefore for these NCP, we assume NCP capacity to be equal to the supply, following previous studies (Verhagen et al., 2017; O’Connor et al., 2021).

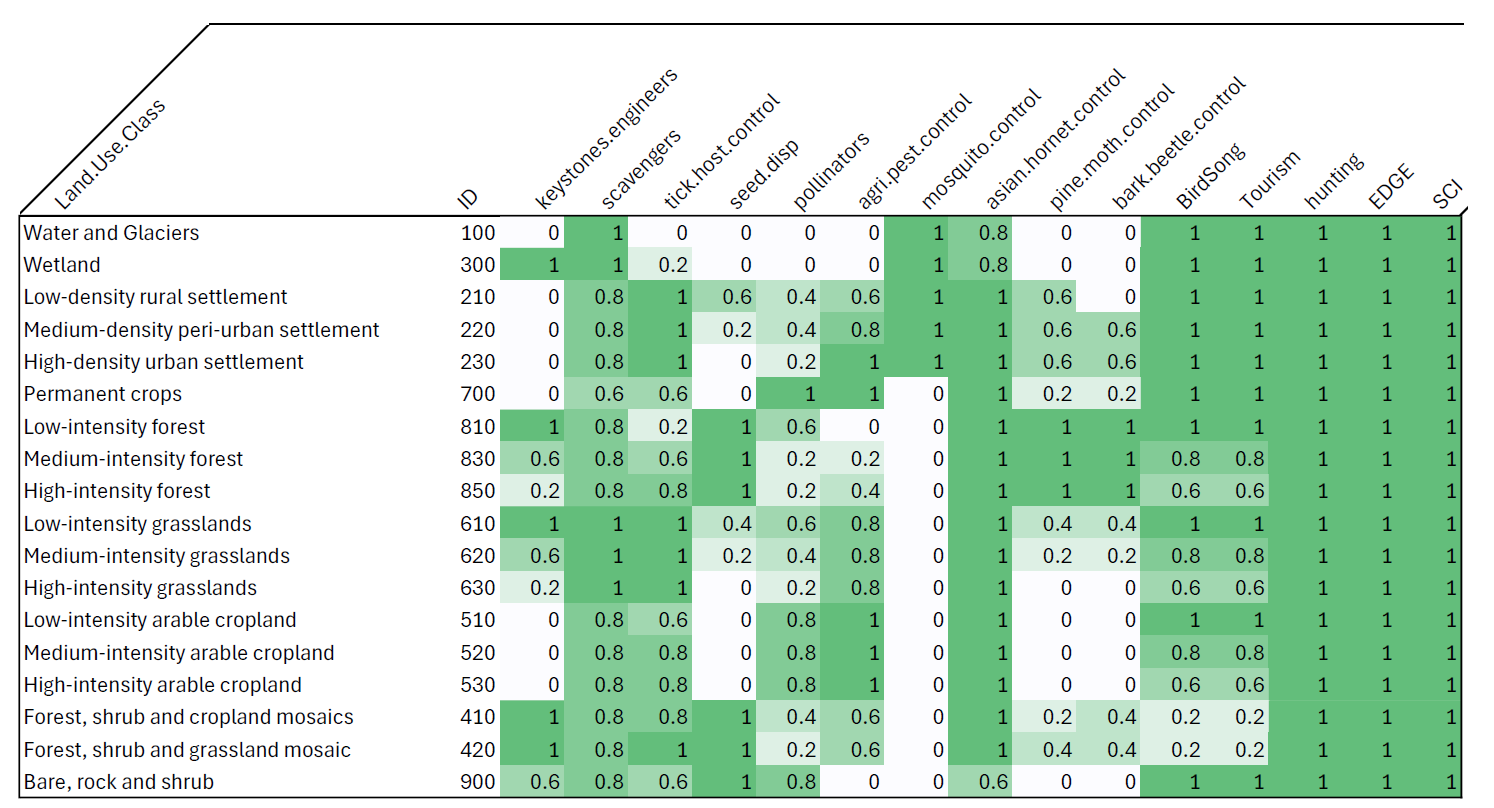

1. Aves : <https://doi.org/10.15468/dl.6ptbvk> ; <https://doi.org/10.15468/dl.6am8ch> ; <https://doi.org/10.15468/dl.6q3zpq> ; <https://doi.org/10.15468/dl.y8nfqw> ; <https://doi.org/10.15468/dl.gdta5n> ; <https://doi.org/10.15468/dl.tn2b3w> ; <https://doi.org/10.15468/dl.2bhy85> ;

   Amphibia/Mammalia/Reptilia: <https://doi.org/10.15468/dl.rxfzz3>
